# NOTCH-YAP1/TEAD-DNMT1 axis regulates hepatocyte reprogramming into intrahepatic cholangiocarcinoma

**DOI:** 10.1101/2020.12.03.410993

**Authors:** Shikai Hu, Laura Molina, Junyan Tao, Silvia Liu, Mohammed Hassan, Sucha Singh, Minakshi Poddar, Aaron Bell, Daniela Sia, Michael Oertel, Reben Raeman, Kari Nejak-Bowen, Aatur Singhi, Jianhua Luo, Satdarshan P. Monga, Sungjin Ko

## Abstract

Intrahepatic cholangiocarcinoma (ICC), a disease of poor prognosis, has increased in incidence. It is challenging to treat due to intra- and inter-tumoral heterogeneity, which in part is attributed to diverse cellular origin. Indeed, co-expression of AKT and NICD in hepatocytes (HCs) yielded ICC, with similarity to proliferative, Notch-activated, and stem cell-like subclasses of clinical ICC. NICD regulated SOX9 and YAP1 during ICC development. *Yap1* deletion *or TEAD* inhibition impaired HC-to-biliary epithelial cell (BEC) reprogramming and ICC proliferation; *Sox9* loss repressed tumor growth; and *Yap1-Sox9* combined loss abolished ICC development in AKT-NICD model. DNMT1 was discovered as a novel downstream effector of YAP1-TEAD complex that directed HC-to-BEC/ICC fate-switch. DNMT1 loss prevented Notch-dependent HC-to-ICC development, and DNMT1 re-expression restored ICC development following TEAD repression. Coexpression of DNMT1 with AKT was sufficient to induce hepatic tumor development including ICC. Thus, we have identified a novel NOTCH-YAP1/TEAD-DNMT1 axis essential for HC-driven ICC development.

**SIGNIFICANCE:** We evaluated the clinical relevance of hepatocyte-driven ICC model and revealed critical but distinct roles of YAP1 and SOX9 in AKT-NICD-driven hepatocyte-derived ICC. We also identified NOTCH-YAP1/TEAD-DNMT1 axis as a critical driver for hepatocyte-to-ICC reprogramming, which might have biological and therapeutic implications in ICC subsets.

## INTRODUCTION

Intrahepatic cholangiocarcinoma (ICC) has steadily risen in the US, while the five-year survival rate of ICC remains under 10% (1). Although FGFR inhibitor was recently approved as first targeted therapy (2) and IDH1/2 inhibitors are in phase 3 clinical trial (2,3), the impact of these inhibitors is limited since only ~15% of patients harbor these aberrations and long-term survival benefits are only modest (4–6). Thus, there is a dire need for better understanding of the diverse molecular mechanisms that drive ICC, in order to develop targeted therapies and identify biomarkers for precision medicine.

ICC is traditionally considered to originate from cholangiocytes (biliary epithelial cells; BECs), as tumor cells exhibit luminal structures and express BEC-specific markers such as SOX9 and Cytokeratin (CK)-19. However, a subset of human ICC has also been theorized to originate from hepatocytes (HCs), based on the frequent detection of human ICC in the pericentral zone of the liver lobule (which is anatomically distinct from the native biliary structure), the high prevalence of mixed HCC-ICC in human patients, and the well-proven occurrence of HC-to-BEC differentiation in various murine models of chronic cholestasis by lineage tracing systems (4,7–11). This theory has been supported by several studies demonstrating a rapid onset of HC-derived murine ICC by either thioacetamide (TAA) administration (12) or by delivery of combinations of some oncogenes such as myristoylated-*Akt* (referred hereon as Akt) and Notch intracellular domain (NICD), *Akt-Yap, Akt-Fbxw7□F or KRAS-sh-p53* into HCs using the sleeping beauty transposon-transposase and hydrodynamic tail vein injection (SB-HDTVI) technique (8,9,11,13,14).

*Sox9* and *Yap* are well known biliary-specific Notch target genes or regulators of Notch signaling, and hence critical in BEC maturation and bile duct morphogenesis in murine development (15). Expression of YAP or SOX9 is prevalent in human ICC and positively correlates with clinical grade (16,17). Despite these observations, the exact significance, pathologic roles and mechanisms by which these factors may lead to ICC, have remained poorly understood.

Herein, we reveal that *Akt-NICD*-driven HC-derived ICC model shows high genetic similarity to a subset of human ICC cases, and provide novel pathologic and mechanistic insights into how SOX9 and YAP/TEAD-DNMT1 axis contribute to the HC-driven-cholangiocarcinogenesis. We elucidate distinct yet cooperative interactions among various reprogramming molecules that drive HC transformation to yield ICC, underscoring the disparate cellular origin of these tumors, which may have both biological and therapeutic implications.

## RESULTS

### p-AKT, and NICD effectors YAP and SOX9, are activated in pre-malignant liver diseases and also prevalent in human cholangiocarcinoma

Given that SB-HDTVI induces HC-driven ICC in AKT-NICD and AKT-YAP mouse models (8,11), we first explored if any of these ICC driver genes are actually evident in HCs of patients with chronic liver diseases that are known ICC risk factors. By immunohistochemistry (IHC), we examined any evidence of active AKT signaling (phospho-Ser473-AKT or p-AKT) and nuclear localization of SOX9 and YAP, both known targets of NICD (15,18), in liver samples from patients with non-alcoholic steatohepatitis (NASH) and primary sclerosing cholangitis (PSC), known risk factors for CCA (10,19). We identified significant aberrant induction of cytoplasmic and nuclear p-AKT (p=0.024), nuclear SOX9 (p=0.0008) or nuclear YAP (p=0.047) in subsets of HCs in all patients (n=20), whereas these markers were rarely detected in hepatocytes of healthy controls (Fig.1A, Fig.S1A and S1B, Table S1). Further, 35% of cases showed some degree of upregulation of all three markers, while 55% showed upregulation of at least 2 of the 3 markers in a subset of HCs (Fig.1B and Fig.S1B-E). These results may imply that in chronic liver injuries in patients, specifically those associated with a risk of ICC development, there is evidence of ectopic activation of ICC drivers (AKT, SOX9 and YAP) in HCs, which may prime these cells for subsequent transformation with additional hits over time.

**Figure 1.**
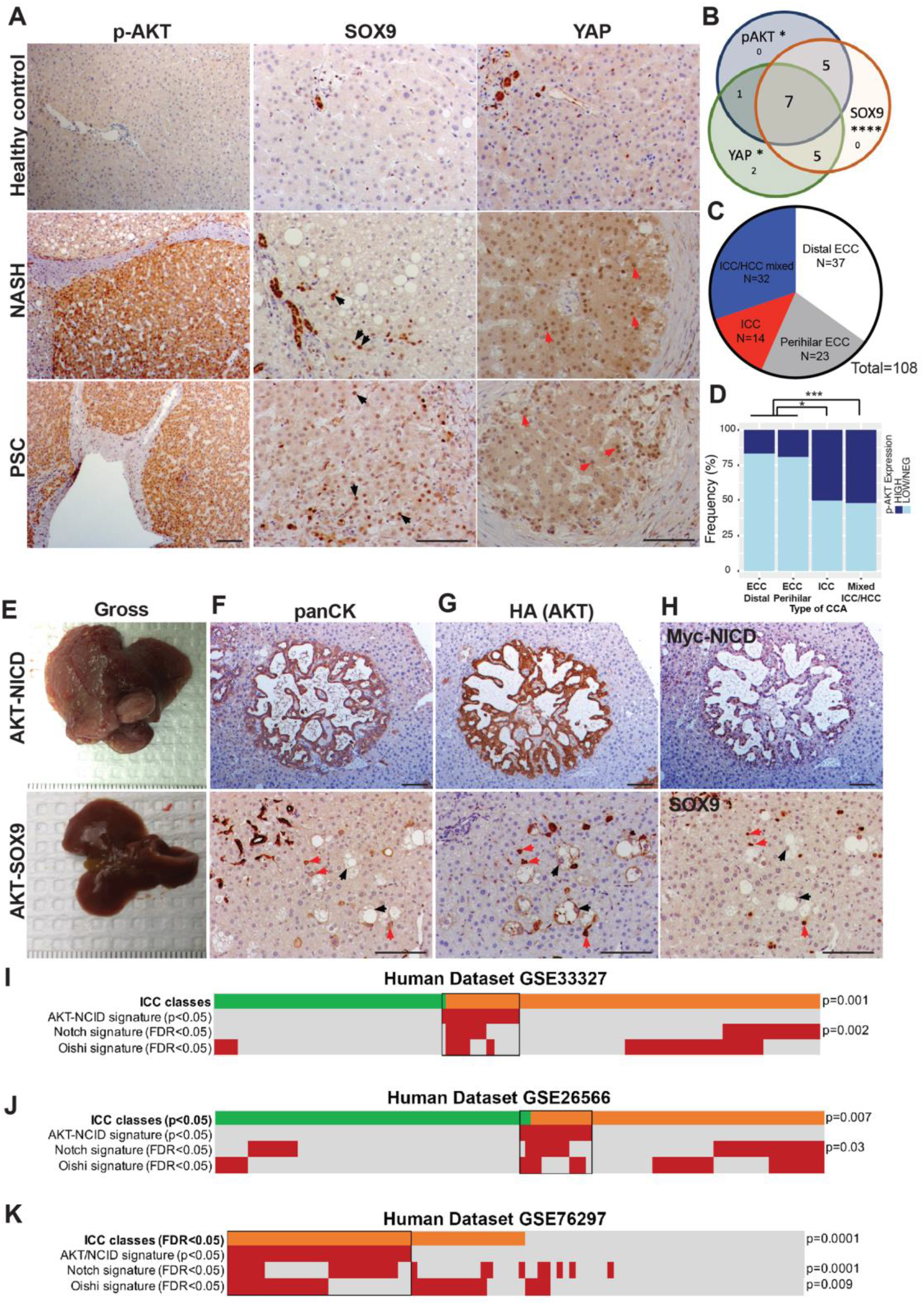
Clinical relevance of AKT-NICD-mediated HC-driven murine ICC. **(A)** Representative IHC images of liver section from patients with NASH and PSC showing increased p-AKT, SOX9 and YAP expression as compared to healthy liver. Black arrows point to YAP^+^ hepatocytes; red arrows to Sox9^+^ hepatocytes; white arrows to p-AKT^+^hepatocytes. (**B**) Venn diagram showing the overlap of patient samples with NASH and PSC that exhibited aberrant induction of either p-AKT, SOX9, or YAP levels in HCs specifically. **(C)** Distribution of subtypes of CCA on TMA containing 108 patients analyzed for AKT and NICD targets by IHC. **(D)** Stratifying p-AKT IHC staining by CCA subtypes shows an enrichment of p-AKT-HIGH staining in ICC and mixed ICC-HCC tumors, while extrahepatic CCA tended to have LOW or negative p-AKT. **(E)**Gross images of livers from AKT-NICD (upper panel) and AKT-Sox9 injected mice (lower panel) at 5 weeks post-HDTVI of respective SB plasmids along with SB transposase. **(F)** Representative IHC staining for panCK depicts presence of ICC in AKT-NICD model (upper panel) and lack of any tumors with positive staining in normal ducts. Red arrows indicate HA-AKT and SOX9-transfected hepatocytes and black arrows point to HA-AKT-transfected but SOX9-negative foamy hepatocytes in the AKT-SOX9 model (lower panel). **(G)** Representative IHC staining for HA-tag to identify myr-*Akt* in serial sections (to F) depicts its presence in ICC in AKT-NICD model (upper panel). Red arrows indicate HA-AKT and SOX9-transfected hepatocytes and black arrows point to HA-AKT-transfected but SOX9-negative foamy hepatocytes in AKT-SOX9 model (lower panel). (**H**) Representative IHC staining on serial sections (to F and G) to identify MYC tag to identify NICD (upper panel) and SOX9 (lower panel) at 5-weeks post SB-HDTVI in AKT-NICD and AKT-SOX9 livers. Nearest Template Prediction (NTP) analysis of three human whole-tumor gene
expression datasets GSE33327 **(I)**, GSE26566 **(J)**and GSE76297 **(K)**using the differentially expressed gene signature comparing wild-type mouse liver and AKT+NICD-injected liver generated in this study. Each human sample was assigned to one of the two ICC molecular classes previously described by *Sia D et al Gastroenterology 2013;*Inflammation class is indicated in green, Proliferation class is indicated in orange. In the heatmap, each column represents a patient and each row represents a different signature; positive prediction of signatures as calculated by NTP is indicated in *red* and absence in *gray*. Notch signature used herein is derived from *Villanueva et al, Gastroenterology 2012* and Oishi signature, which captures a hepatic stem-cell like group of ICC patients, was derived from *Oishi et al, Hepatology 2012*. p values that show significant correlation are indicated to the right of the NTP analysis. Scale bars:100 μm; *p<0.05; ***p<0.001; ****p<0.0001.

Next, by IHC, we examined the localization of p-AKT, SOX9 and YAP in CCA using 2 tissue microarrays (TMAs), containing 1-4 tissue samples each from 108 CCA patients, including 14 ICCs, 32 intrahepatic mixed HCC-ICC samples, and 61 extrahepatic CCAs (UPMC, Tables S2 and Fig.1C). IHC staining for each marker was scored as described in the Methods (Fig. S1F). Of the 108 intact readable samples, 103 (95%) showed high nuclear YAP and 102 (94%) showed prominent nuclear SOX9 localization, consistent with previous reports (11,16,17). Of the 102 intact readable samples, 32 showed strong cytoplasmic p-AKT immunostaining. Interestingly, we detected a strong enrichment of p-AKT expression in ICC (7/14, 50%, p=0.024) and intrahepatic mixed HCC-ICC samples (15/31, 48%, p=0.006) as compared to extrahepatic CC (10/58, 17%), by the Fisher’s exact test with pairwise comparison (Fig.1D). This suggests that activation of AKT may contribute specifically to the pathogenesis of ICC, and lends support to our hypothesis that ectopic activation of AKT in HCs may be a risk factor for HC-derived ICC. This data further supports the clinical relevance of studying SB-HDTVI-driven tumorigenesis in AKT-NICD and AKT-YAP murine models.

### Myristoylated-AKT cooperates with NICD but not SOX9 in inducing hepatic tumors

Based on preclinical and clinical observations, we next established SB-HDTVI model to explore the role of NICD effectors YAP and SOX9 with AKT activation in hepatic tumorigenesis. As reported previously as well, we observe rapid development and growth of HC-derived ICC 4-5 weeks after HDTVI of myr-*Akt* (HA tag) and *NICD* (Myc tag) in the AKT-NICD model (Fig.1E-H) (8,11). IHC confirmed the ICCs to be positive for biliary marker pan-cytokeratin (panCK), HA tag (AKT), MYC tag (NICD), and SOX9 where expected (Fig.1F-H). Given the critical role for SOX9 downstream of Notch signaling in HC-to-BEC conversion in liver injury (18,20), we next generated AKT-SOX9 model by HDTVI of myr-*Akt* (HA tag) and *Sox9*. Interestingly, no tumors were observed at 5-weeks post-injection, either macroscopically or microscopically (Fig.1E-H) and even up to 4 months (data not shown). Histologically, cells transduced with *Akt* and *Sox9* only weakly expressed panCK and exhibited an intermediate morphology between HCs and BECs, but they remained as single cells without any evidence of clonal expansion to form tumors at 5-weeks (red arrows in Fig.1F-H). Our results suggest that SOX9 is not sufficient to drive HC-derived ICC downstream of Notch signaling, despite concurrent AKT activation.

### Transcriptome analysis of HC-derived ICC in AKT-NICD model reveals significant similarity to a subset of human ICCs

To address the clinical relevance of the ICCs observed in the AKT-NICD model, we performed RNA-sequencing (RNA-Seq) analysis. When comparing the WT and AKT-NICD livers, 641 genes were upregulated and 241 genes were down-regulated, by FDR=5% and absolute log2 fold change greater than 1 (Fig.S2A). To interpret biological functions of these differentially expressed genes (DEGs), pathway enrichment analysis was performed by Ingenuity Pathway Analysis. Thirty-six pathways were significantly enriched in the AKT-NICD livers. To determine if the mouse model mimics a subtype of patient ICC, a human study GSE33327 of ICC was analyzed (21), using similar pipeline. When comparing 6 control and 149 ICC categorized as inflammatory (n=57) and proliferative (n=92), 590 upregulated and 781 down-regulated genes were enriched in 154 pathways. Seven pathways were commonly altered in mouse and human tumors, including PKC-theta signaling in T lymphocytes pathway, maturity onset diabetes of young (MODY) signaling pathway, histamine biosynthesis pathway, pentose phosphate pathway, netrin signaling pathway, nicotine degradation III pathway and GP6 signaling pathway. Likewise, another human study (GSE26566) consisting of 104 CCA and 59 surrounding liver samples, was assessed (22). Using FDR of 1% absolute log2 fold change of 3 or greater, 1145 upregulated and 1085 down-regulated genes were enriched in 179 pathways, were identified. Fourteen pathways were commonly altered in mouse and human tumors including Maturity Onset Diabetes of Young (MODY) Signaling and GP6 Signaling Pathway.

To directly compare mouse and human study, top DEGs from the AKT-NICD model were converted to human homologous genes by the Mouse Genome Database (23). Top DEGs were then applied to human study GSE33327 and 44 top genes are shown (Fig.S2B). These genes could clearly separate the human normal (orange bar), inflammation (light green bar) and proliferation (pink bar) ICC subgroups in patients very well. Similarly, DEGs from the AKT-NICD mouse model were also applied to GSE26566. Top 242 of the 522 genes with abs(log_2_FC)>1 and FDR<0.05 were selected. These genes could clearly stratify human normal surrounding liver (orange bar) and CCA (light green bar) (Fig.S2C). DEGs from the AKT-NICD mouse model were also applied to another human study GSE76297, composed of CCA patients from Thailand (24). Top 53 DEGs could also separate the human non-tumor samples (orange bar) and CCA samples (lightgreen bar) in this dataset are shown (Fig.S2D).

Next, Nearest Template Prediction (NTP) analysis was performed using the AKT-NICD signature. In GSE33327, a small percentage of human patients (13%, 19/149) were positively predicted by the AKT-NICD signature (Fig.1I, orange bar). Patients molecularly similar to the AKT-NICD murine model belonged mostly to the “proliferation class” (18/92 vs 1/57, p=0.0009), as defined previously (21). In the GSE26566 dataset, around the same percentage (12%; 13/120) of human ICC patients were predicted to be positive by the AKT-NICD signature and 11 of them belonged also to the proliferation class (11/53 vs 2/57, p=0.007) (Fig. 1J, orange bar). In the Thailand dataset GSE76297, the AKT-NICD signature predicted the proliferation class of ICC (30%; 27/91), while no inflammation class was observed in this dataset (Fig.1K). In addition, in all three datasets, the group of human patients resembling the AKT-NICD murine model were enriched in NOTCH signaling (p<0.05) (25) and in 1/3 studies in the previously reported stem cell like signature of human ICC (26), (Fig.1I-K). Altogether, the AKT-NICD model is an excellent representation of a subset of human ICC and suggests possible HC origin of ICC in this subset of cases.

### Conditional *Yap* or *Sox9* deletion significantly delays cholangiocarcinogenesis in the AKT-NICD model

Given the high nuclear levels of YAP and SOX9 in human CCA and the well-known interaction of NOTCH, SOX9 and YAP in the biliary compartment (4,10,11,16–18,27), we next investigated the cell autonomous roles of YAP and SOX9 in the AKT-NICD model. In addition to AKT-NICD, we co-delivered either *pCMV-Cre* into *Yap^flox/flox^* or *Sox9^flox/flox^* mice to delete *Yap* or *Sox9* in transduced HCs, respectively (labeled as *Yap* KO or *Sox9* KO) (Fig.2A). As controls, *pCMV-Empty* was injected into *Yap^flox/flox^* or *Sox9^flox/flox^* mice and for analysis, both groups were combined and called wildtype (WT). All WT mice developed a lethal burden of ICC and had to be euthanized by 5-6-weeks post SB-HDTVI (Fig.2B-F). In contrast, only one of six *Sox9* KO mice became sick due to tumor burden requiring euthanasia at 5 weeks, while the remaining five *Sox9* KO mice showed no signs of illness and survived until sacrifice at 38 days for comparison to WT. All AKT-NICD *Yap* KO mice were healthy at 6 weeks, when they were also sacrificed for comparison to WT (Fig.2A and 2B). Grossly, AKT-NICD *Yap* or *Sox9* KO had only rare or no tumor and significantly lower liver weight-to body weight ratio (LW/BW) compared to widespread gross disease in WT (Fig.2C-F). Histology also verified presence of small and/or rare microscopic panCK-positive tumor foci in *Yap* or *Sox9* KO deleted livers (Fig.2E). In fact, panCK-positive area compared in both KO was significantly less than WT (Fig.2F). Altogether, conditional elimination of either *Yap* or *Sox9* significantly diminished HC-derived ICC in the AKT-NICD model.

**Figure 2.**
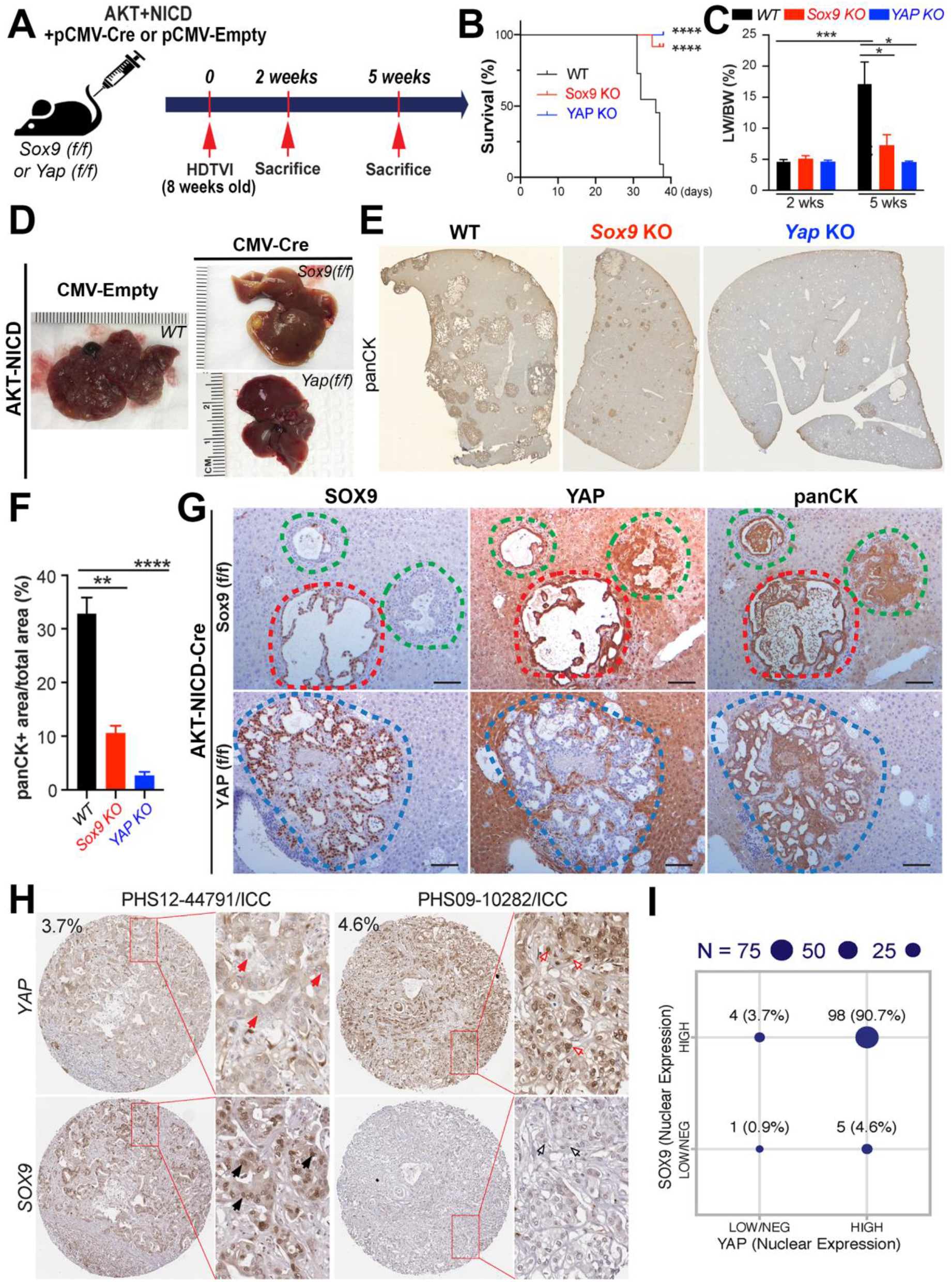
Tumor-specific Sox9 or Yap deletion reduces AKT-NICD-induced hepatocyte-derived ICC development. **(A)** Experimental design illustrating plasmids used for HDTVI, mice used in study and time-points analyzed. **(B)** Kaplan–Meier survival curve showing improved survival of *Sox9* KO and *Yap* KO as compared to WT upon establishment of AKT-NICD model. **(C)** LW/BW ratio depicts comparable low tumor burden in AKT-NICD *Sox9* KO, Yap KO and WT mice at 2 weeks but notable increase in tumor burden in WT at 5 weeks, which was significantly lower in *Sox9* KO and *Yap* KO mice. **(D)** Representative gross images from AKT-NICD WT show multiple and large tumors at 5 weeks, with only occasional small tumor seen in *Sox* KO and almost no gross tumor in *Yap* KO at the same time-point. **(E)** Representative tiled image of panCK staining showing typical papillary and cystic pattern in AKT-NICD WT mice with *Sox9* KO showing fewer and smaller foci and *Yap* KO showing even fewer lesions at 5 weeks. **(F)** Quantification of panCK IHC verifies significantly reduced staining in *Sox9* KO and *Yap* KO as compared to WT at 5 weeks. **(G)** IHC staining for SOX9, YAP and panCK showing ICC positive for all markers (red dashed lines) as well as SOX9^−^ but YAP and panCK positive ICC (green dashed lines) in AKT-NICD *Sox9* KO at 5 weeks. Likewise, occasional tumor observed in AKT-NICD *Yap* KO was YAP^−^ but positive for SOX9 and panCK (blue dashed lines). (**H**) TMA sections from CCA patients were stained for SOX9 and YAP. Enlarged images are shown for better appreciation of nuclear localization of SOX9 and YAP. Red arrows point to nuclear YAP^−^ cells; black arrows point to nuclear SOX9^+^ cells; red empty arrows point to nuclear YAP^+^ cells; black empty arrows point to nuclear SOX9^−^ cells. **(I)** Correlation of YAP and SOX9 nuclear staining in human CCA samples from TMA shows majority of tumors to be positive for both markers, but a small fraction to be SOX9^+^ and YAP^−^ (or low) or SOX9^−^ (or low) and YAP^+^ (representative IHC images shown in (H). Scale bars: 100 μm; error bar: standard error of the mean; *p<0.05; **p<0.01; ***, p<0.001; ****p<0.0001.

Since we were delivering 3 plasmids (*Akt, NICD, pCMV-Cre/pCMV-Empty*) concurrently, we next asked if some tumor nodules in the *Yap* KO and *Sox9* KO livers resulted from HCs which may have failed to take up *pCMV-Cre*-plasmid and hence escape deletion of *Yap* or *Sox9*, respectively. Indeed, in the livers from *pCMV-Cre*-injected mice (both *Sox9^flox/flox^* or *Yap^flox/flox^*), although tumor number and size were greatly reduced compared to the WT, both SOX9 and YAP positive tumors were detected in the *Sox9* KO and *Yap* KO (Fig.2G, red dashed line in *Sox9* KO and not shown for *Yap* KO). Intriguingly though, a small number of ICC in *Sox9* KO were SOX9-negative and positive YAP-positive (Fig.2G, green dashed lines). Similarly, a small number of ICC in *Yap* KO were YAP-negative and SOX9-positive (Fig.2G, blue dashed lines). Lastly, to address clinical relevance of singly positive ICC for SOX9 or YAP, we examined patient TMA and identified YAP^+^;SOX9^−^ (4.6%) or SOX9^+^;YAP^−^ (3.7%) CCA (Fig. 2H and 2I). These observations support the clinical relevance of murine findings and suggest unique subsets of ICC that may be driven singly by YAP or SOX9, likely with distinct molecular signatures.

### Distinct and overlapping roles of YAP and SOX9 in AKT-NICD driven cholangiocarcinogenesis in regulating reprogramming, survival and proliferation

We next sought to address specific functions of SOX9 and YAP in the HC-derived ICC development. To investigate, we eliminated either *Sox9* or *Yap* using *pCMV-Cre* (as shown in Fig.2A) co-injection along with *Akt* and *NICD* and examined ICC after 5 weeks for proliferation by immunofluorescence for PCNA, and cell viability by IHC for TUNEL (Fig.3A). Interestingly, tumors with absence of YAP or SOX9, both exhibited a considerable and significant decrease in the numbers of PCNA^+^ cells than WT (Fig.3B and 3C). Additionally, cell death was notably and significantly increased in the absence of SOX9 but not YAP, as compared to the WT (Fig.3D and 3E).

**Figure 3.**
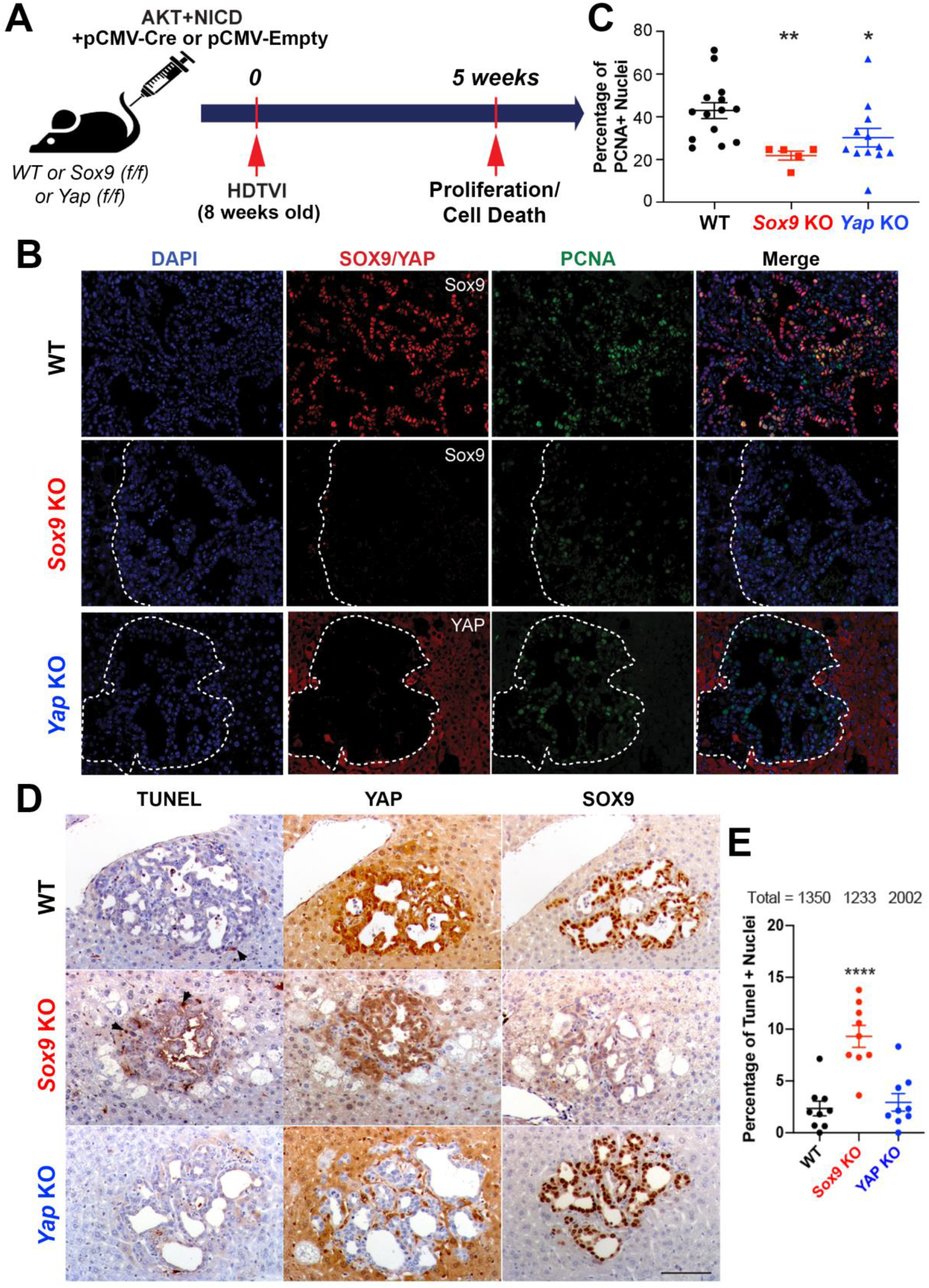
Distinct and overlapping roles of Sox9 and Yap in AKT-NICD-induced hepatocyte-derived ICC. (**A**) Experimental design illustrating plasmids used for HDTVI, mice used in study and time-points analyzed. (**B**) Representative IF for SOX9 (red), YAP (red), PCNA (green) and DAPI (blue) in liver sections from 5 weeks AKT-NICD WT, Sox9 KO or Yap KO mice. **(C)** The percentage of PCNA-positive nuclei normalized to total tumor cell nuclei in representative images shown in B, demonstrate significant decreased in proliferation in Sox9 KO and Yap KO as compared to WT. **(D)** IHC for TUNEL to detect non-viable tumor cells, shows higher cell death in Sox9 KO only while minimal and comparable cell death was evident in WT and Yap KO in AKT-NICD model at 5-week time-point. **(E)**The percentage of TUNEL-positive nuclei normalized to total tumor cell nuclei in representative images shown in D, demonstrate significant increase in cell death in Sox9 KO as compared to WT or Yap KO. Scale bars:100 μm; error bar: standard error of the mean; *p<0.05; **p<0.01; ****p<0.0001.

Given the roles of YAP and SOX9 in biliary fate determination (27), we next investigated HC-to-BEC reprogramming in the *Akt* and *NICD* transduced HCs in WT, *Sox9* KO and *Yap* KO, 2 weeks post HDTVI in the AKT-NICD model (Fig.4A). At this early stage, *Akt* and *NICD*-transduced cells in WT livers begin to display biliary morphology and positivity for SOX9, YAP and panCK, and lose expression of HNF4α as shown by IHC in serial sections (Fig.4B, red dashed line). These observations were also substantiated by immunofluorescence (Fig.4C). However, upon *Yap*-deletion the *Akt-NICD* transduced cells continue to exhibit HC morphology and remain HNF4α-positive as seen by IHC in serial sections as well as by immunofluorescence, indicating a defective HC-to-BEC trans-differentiation (Fig.4B, blue dashed line). Interestingly, a subset of *Yap*-deleted cells co-express SOX9 and HNF4α (Fig.4C, white arrows), which may represent an intermediate cell type with incomplete HC-to-BEC reprogramming. Importantly, this intermediate cell population was not detected at 2 weeks in the AKT-NICD WT mice. Surprisingly, *Sox9*-deleted *Akt-NICD* transduced cells showed similar early stage biliary morphology as WT with positive biliary markers (YAP and panCK) and absence of HNF4α, by both IHC on serial sections (Fig.4B, green dashed line) and immunofluorescence (Fig.4C). Altogether, these observations show both unique and overlapping but temporal roles of these HC reprogramming factors and explain the observed repressive effects of *Sox9* or *Yap* deletion in the HC-derived cholangiocarcinogenesis.

**Figure 4.**
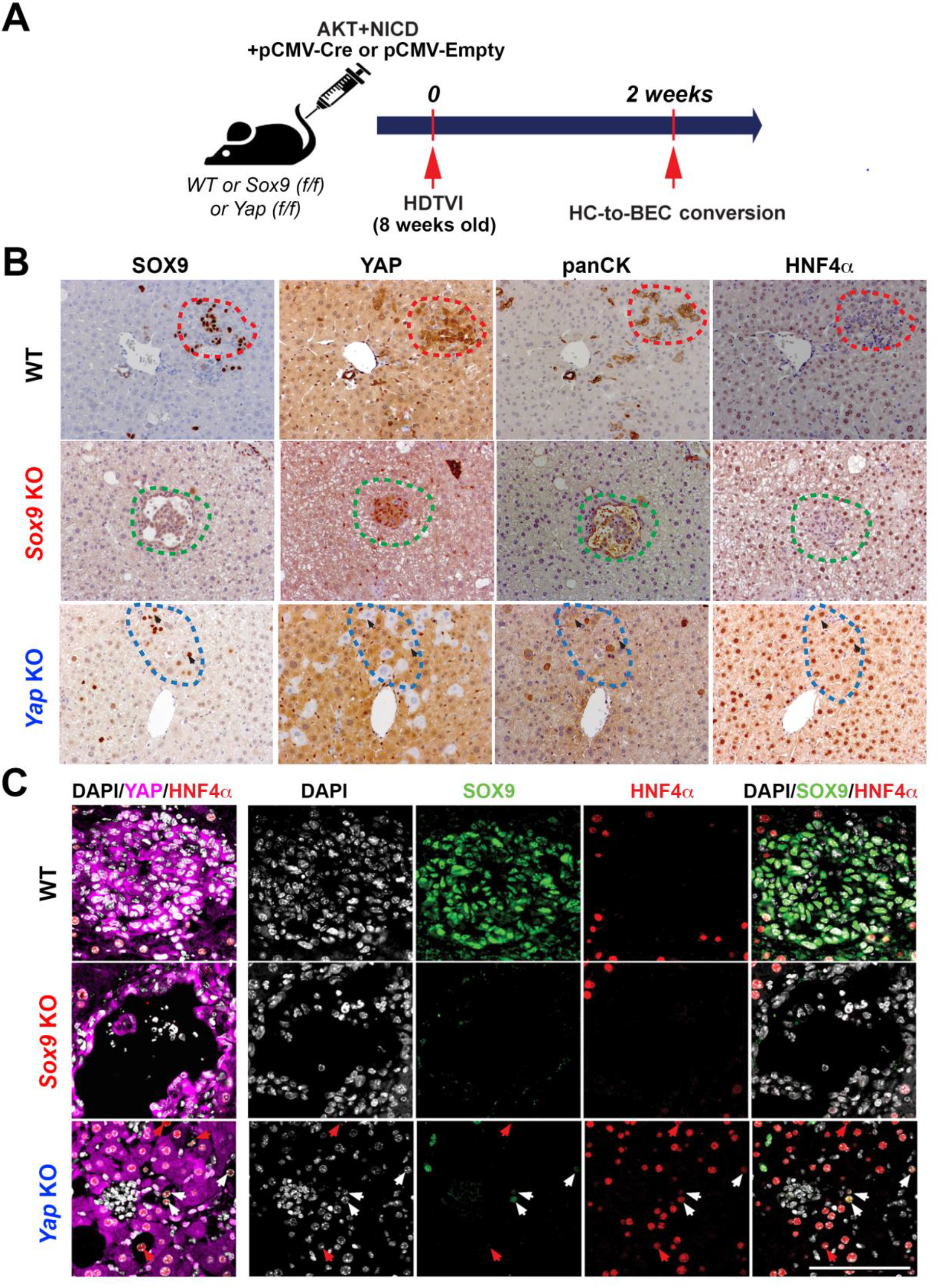
Role of Yap but not Sox9 in hepatocyte-to-biliary reprogramming in AKT-NICD-induced cholangiocarcinogenesis. (**A**) Experimental design illustrating plasmids used for HDTVI, mice used in study and time-points analyzed. (**B**) Representative IHC staining of WT livers 2 weeks after AKT-NICD injection showing earliest transformed hepatocytes losing staining for HNF4α and acquire biliary morphology and expression of SOX9, YAP and panCK (red dashed lines). Sox9 KO livers showing Sox9 deleted AKT-NICD-transfected cells with intact HC-to-biliary reprogramming at the 2-week time point with loss of HNF4α and gain of YAP and panCK despite SOX9 loss (green dashed lines). Yap KO livers showing Yap deleted AKT-NICD-transfected cells with imperfect HC-to-biliary reprogramming at the 2-week time point with continued HNF4α staining and some staining for SOX9 and panCK in YAP-negative HCs (blue dashed lines). **(C)** Confocal images of IF staining of WT, Sox9 KO and Yap KO livers at 2 weeks after AKT-NICD injection verify IHC results in B. Red arrows point to YAP-negative, HNF4α-positive and Sox9-negative cells and white arrows to YAP-negative, HNF4α-positive and Sox9-positive cells. Scale bars:100 μm

### Simultaneous *Yap* and *Sox9* loss prevents AKT-NICD-driven ICC

Based on the identification of SOX9 or YAP single positive AN-ICC (Fig.2G), we next investigated the impact of concurrently eliminating the expression of *Sox9* and *Yap* in the AKT-NICD model. We injected *NICD* and *pCMV-Cre* along with *myrAkt-shYap* plasmids into the *Sox9^flox/flox^* mice to achieve tumor-specific dual knockout (dKO) of *Sox9* and *Yap* in the transduced HCs (Fig.5A). As controls, we injected *NICD, pCMV-Empty* and *myrAkt-sh-Luciferase* (*sh-Luc*) plasmid into the *Sox9^flox/flox^* mice to generate mouse wild-type for dual genes (dWT) (Fig.5A). Consistent with previous results, AKT-NICD dWT mice developed lethal ICC requiring euthanasia around 5-weeks post-HDTVI whereas dKO were asymptomatic at 5 weeks and 3-months post-HDTVI, showing significant differences in survival (Fig.5B). LW/BW assessment showed significantly lower tumor burden in dKO at both 5 weeks and 3 months as compared to dWT at 5 weeks (Fig.5C). Gross examination of the livers verified these observations at all time points (Fig.5D). Histological analysis at 5 weeks by IHC for HA tag (AKT), MYC tag (NICD) and panCK, confirmed that AKT-NICD dWT livers develop ICC composed of transdifferentiated HC, whereas dKO livers showed complete absence of any tumors at 5 weeks (Fig.5E). These data clearly suggest that dual suppression of *Yap* and *Sox9* is sufficient to completely abrogate *Akt-NICD*-dependent HC-derived ICC development (Fig.5C-E).

**Figure 5.**
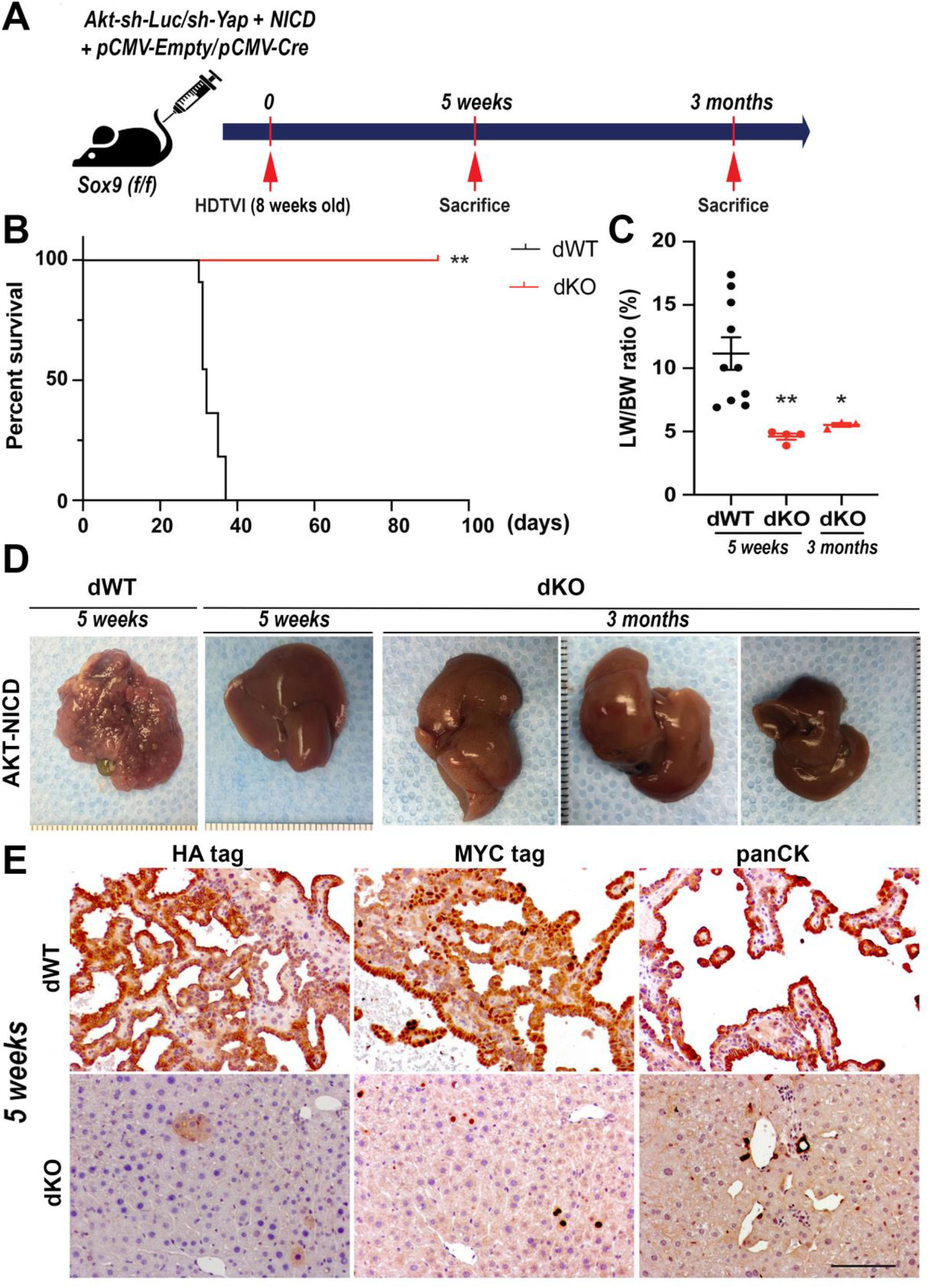
Simultaneous *Yap* and *Sox9* elimination completely abrogates AKT-NICD driven HC-derived ICC. (**A**) Experimental design illustrating plasmids used for HDTVI, mice used in study and time-points analyzed. **(B)** Kaplan–Meier survival curve showing significantly improved survival of *Sox9* and *Yap* double KO (dKO) as compared to WT upon establishment of the AKT-NICD model. **(C)** LW/BW ratio depicts significantly lower tumor burden in AKT-NICD dKO at both 5 weeks and 3 months post-injection as compared to WT mice at 5 weeks. **(D)** Representative gross images from AKT-NICD WT mice showing multiple large tumors at 5 weeks, with absence of any gross tumors in dKO at either 5-week or 3-month time point. **(E)** Representative IHC images for HA-tag, MYC-tag, and panCK showing dramatic tumor development in AKT-NICD WT livers and normal histology in dKO at 5 weeks after injection. Scale bars:100 μm; error bar: standard error of the mean; *p<0.05; **p<0.01.

### DNMT1 is required for NOTCH-YAP-driven HC-to-BEC/ICC fate-switch

Given the indispensable roles for YAP in Notch-dependent HC-to-BEC conversion (Fig.4), we next sought to delineate the molecular mechanism by which YAP regulates HC-to-BEC and eventually ICC fate-switch downstream of the Notch signaling. A recent study revealed *Nuak2* as a key downstream factor for YAP-driven HCC development by ChIP-seq (28). However, *Nuak2* expression did not change in *Akt-NICD*-driven ICC from our RNA-seq data suggesting that *YAP* regulates unique gene/s that drive HC-driven cholangiocarcinogenesis. To explore possible YAP target genes in hepatocyte, we first re-analyzed public ChIP-seq data analyzing HC-specific YAP over-expression (28) since *Akt-NICD* transduction also induced strong YAP activation, critical for HC-to-BEC reprogramming (Fig.2–4). Through this approach, we identified 393 potential TEAD4 target genes in YAP-active HC. Among TEAD4-bound genes in YAP-activated HC, we next focused on a master epigenetic regulator DNA Methyl Transferase-1 encoded by *Dnmt1* (Fig.S3A). In fact, DNMT1 has been previously show to play an important role in cell fate switches including HC-to-BEC conversion (29–32). We confirmed existence of TEAD2/3/4-specific binding motif in the *Dnmt1* promoter (Fig.S3A-B). Based on these observations, we hypothesized that YAP/TEAD complex upregulates *Dnmt1* transcription, and subsequently DNMT1 represses epigenetic targets which may be inhibiting the conversion of HC into BEC and ICC in the *Akt-NICD* model.

To investigate the roles for DNMT1 in HC-driven cholangiocarcinogenesis, we employed genetic and pharmacologic approaches: 1) FDA-approved DNMT1 specific inhibitor 5-Azacytidine into *Akt-NICD*-ICC model (Fig.6E).; 2) injection of *myrAkt-sh-Dnmt1* plasmid along with NICD to develop *Akt-NICD*-ICC in the setting of *Dnmt1* silencing (Fig.6F). Remarkably, both chemical and genetic DNMT1 inhibition completely blocked Notch-YAP-dependent initial HC-to-BEC conversion of transduced HCs at 2 weeks post injection (Fig.6E, black and red dashed line), similar to what was observed in *Akt-NICD-ICC* after YAP deletion (Fig.6D, blue dashed line). Upon DNMT1 inhibition, the *Akt-NICD*-transduced cells continued to display HC morphology with HNF4αas seen by IHC in serial sections, indicating a block in HC-to-BEC trans-differentiation (Fig.6E, black dashed line and Fig.6F, red dashed line), whereas *Akt-NICD*-transduced cells in control livers converted into BEC/ICC to exhibit typical biliary shape expressing SOX9, YAP and panCK, but negative for HNF4αat this stage (Fig.6B, WT or Fig.6C, *Sox9* KO). Given the similarities in phenotype between YAP-deleted and DNMT1-inhibited *Akt-NICD-ICC* as well as well-defined binding of TEAD4 on *Dnmt1* genomic locus (Fig.S3A), we next examined DNMT1 expression patterns in *Akt-NICD*, *Akt-NICD*-*YAP* KO and *Akt-NICD-Sox9* KO liver, to investigate their interaction in ICC setting. Interestingly, in *Akt-NICD* injected liver, DNMT1 was only detected in NICD^+^;YAP^+^ ICC (Fig.6B, WT or Fig.6C, *Sox9* KO), while NICD^+^;YAP^−^;HNF4α^+^ cells (Fig.6D, blue dash line, impaired HC-to-ICC conversion) were clearly negative for DNMT1 as assessed by IHC, suggesting YAP regulation of DNMT1 in Notch-dependent HC-to-ICC conversion. Also, DNMT1 was not detected in any of HCs transduced with *Sox9* alone (Fig.S5B, red dashed line). Moreover, *Akt* singular transduced foamy cells are DNMT1^−^ (Fig.6C, red arrows), whereas NICD^+^;SOX9^−^ cells strongly express DNMT1 (Fig.6C, *Sox9* KO), indicating that AKT and SOX9 are not involved in Notch-YAP-driven DNMT1 induction in ICC. In addition, given the strong DNMT1 expression and its indispensable role in *Akt-NICD*-dependent HC-to-ICC conversion, we also examined DNMT1 expression in other pre-established, molecularly distinct murine HC-driven ICC models: *Akt+Fbxw7□F, Kras+sh-p53* and TAA-induced ICC (Fig.S4E-G). We observed strong DNMT1 expression in all examined ICC models, suggesting that DNMT1 may play a critical role for HC-to-ICC transformation irrespective of model used.

**Figure 6.**
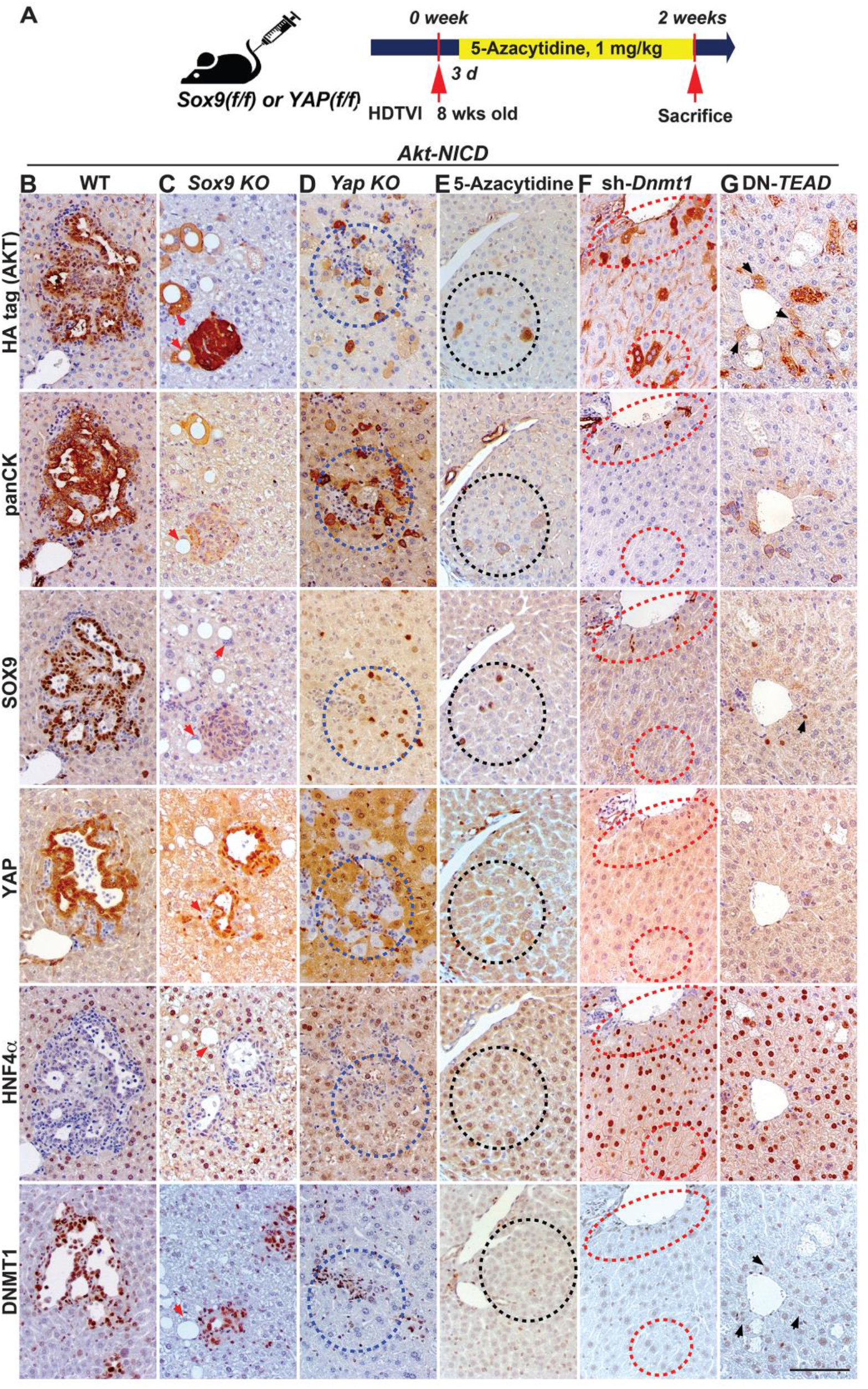
DNMT1 is required for YAP-TEAD-driven HC-to-ICC transformation. **(A)** Experimental design illustrating 5-Azacytidine treatment, plasmids used for HDTVI, mice used in study and time-points analyzed. **(B)** Representative IHC staining of WT or *Sox9* KO livers showing AKT-NICD-transfected cells with intact HC-to-biliary reprogramming with loss of HNF4α and gain of YAP, panCK and DNMT1 regardless of SOX9 expression. *Yap* KO livers showing *Yap*-deleted AKT-NICD-transfected cells with imperfect HC-to-biliary reprogramming with continued HNF4α staining and absent of DNMT1 in YAP^−^ HCs (blue dashed lines). Similarly, 5-Azacytidine-treated, *Dnmt1*-silenced or TEAD-inhibited livers show defective HC-to-BEC reprogramming with continued HNF4α expression and absent of DNMT1 staining in AKT-NICD-transfected cells (black or red dashed lines). Red arrows point *Akt-HA* singular transduced cells. Black arrows point *Akt-HA* transduced cells in *DN-TEAD* liver. Scale bars:100 μm.

To test if DNMT1 is expressed in CCA in patients, which may also suggest HC-to-ICC switch being a more dynamic event in a subset of cases, we assessed 2 TMAs with 108 readable cases. Thirty CCA cases showed strong nuclear DNMT1 (27.8%) (Fig.S3C). All DNMT1^+^ tumors were YAP^+^ but there was no significant enrichment in specific CCA subtypes.

Altogether, these data support a role of DNMT1 in ICC in mice and in patients, especially in ICC originating from HCs.

### NICD-YAP cascade drives HC-to-ICC transformation through the TEAD-dependent DNMT1 induction

Although both YAP or DNMT1 inhibition prevents *Akt-NICD*-driven HC-to-ICC conversion at 2 weeks post-injection, *Yap* singular deletion eventually develops YAP^−^; DNMT1^+^ ICC at 5 weeks post injection (Fig.2G and S4D), suggesting the existence of compensatory mechanisms to replenish transcription of YAP target genes, such as *Dnmt1*. Indeed, it has been recently shown that simultaneous deletion of *Yap* and *Taz* potently represses *Akt-NICD-ICC*, indicating their redundant roles in HC-driven ICC formation under NICD (33). YAP and TAZ proteins are transcriptional coactivators with 47% amino acid sequence identity, encoded by paralogous genes (34). In the nucleus, both proteins require TEAD to bind to DNA and regulate transcription of downstream target genes. Given the TEAD-dependent redundant roles of TAZ and YAP when either is absent (34), we next have asked whether inhibition of TEAD activity using dominant negative (DN)-*TEAD*, which is a truncated mutant lacking DNA binding domain, could permanently block *Akt-NICD*-driven HC-to-BEC conversion and subsequent ICC development, similarly to DNMT1 repression. We injected DN-*Tead* expression plasmid along with *Akt* and *NICD* plasmid to attain tumor-specific TEAD inhibition (Fig.6G). Given that *TEAD1/2/3/4* possess a similar DNA binding motif (35), *DN-TEAD* expression represses transcriptional activity of all TEADs and of YAP-TEAD/TAZ-TEAD complex (35–37). As controls, we injected *NICD, pT3-Ef1 α-Empty* and *AKT* plasmid to develop wildtype *Akt-NICD-ICC* (Fig.6B). Consistent with previous results, *Akt-NICD*-transduced HCs yielded ICC and exhibited biliary morphology expressing SOX9, YAP and panCK, but negative for HNF4αat this stage (Fig.6B). Upon *DN-TEAD* co-delivery, the *Akt-NICD*-transduced cells continued to display HC morphology maintaining HNF4α expression, indicating a block in HC-to-BEC trans-differentiation (Fig.6G, black arrows), similar to what was observed with YAP or DNMT1 inhibition at 2 weeks post injection (Fig.6D-F). Intriguingly, we found that over-expression of *NICD* alone induced fate-switch of HC into SOX9^+^;YAP^+^;panCK^+^;DNMT1^+^;HNF4α^−^ BEC/ICC at 3 weeks post injection, notably fewer than with concomitant AKT activation (Fig.S3G). In contrast, *NICD* and *DN-TEAD* cotransduced cells retained HC morphology with expression of HNF4α but not SOX9, panCK or DNMT1 (Fig.6G), indicating that NOTCH-YAP/TEAD cascade drives HC-to-ICC fate conversion through the regulation of DNMT1 in ICC development.

### DNMT1 or TEAD inhibition permanently abrogate Akt-NICD-driven ICC

Given that DNMT1 or TEAD inhibition completely block NICD-dependent HC-to-ICC conversion at 2 weeks post-HDTVI in the AN model (Fig.6E-G), we next asked if unlike *Yap* deletion, the abrogation in ICC development is permanent. We injected AN along with *DN-TEAD* expression plasmid or treated 5-Azacytidine until sacrifice, and assessed these animals at 5 weeks post-HDTVI (Fig.7A). Consistently, AN-injected control mice developed lethal ICC requiring euthanasia around 5 weeks post-HDTVI while both of DNMT1 or TEAD repressed mice were asymptomatic with comparable LW/BW ratio as healthy mice at 5 weeks post-HDTVI (Fig.7B and C). Grossly, no tumors were observed in any mice with DNMT1 or TEAD repression (Fig.7B and C). Histology verified presence of very rare or no microscopic HA^+^ tumor foci in 5-Azacytidine-treated (Fig.S3E) or *TEAD*-repressed livers (Fig.7D). These data indicate an indispensable role for the TEAD-DNMT1 axis in *Akt-NICD-dependent* HC-driven ICC formation, and also suggested that singular targeting of either is sufficient to eliminate *Akt-NICD-ICC* without any probability of relapse of tumor by activation of a redundant compensatory mechanism, thus overcoming the limitation of tumor suppression by singular targeting of YAP in ICC.

**Figure 7.**
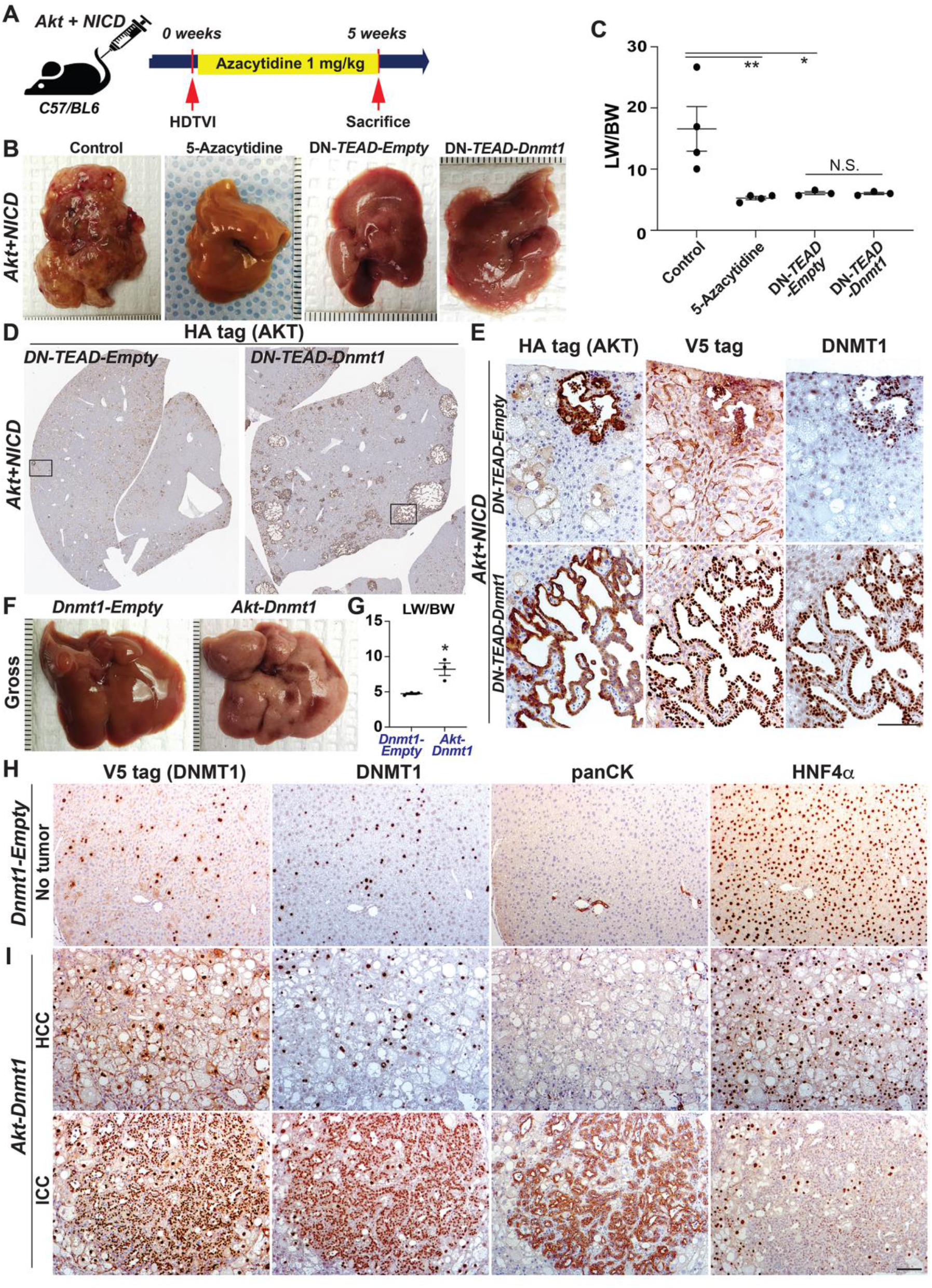
NICD-YAP/TEAD-DNMT1 axis drives HC-to-ICC transformation. **(A)** Experimental design illustrating 5-Azacytidine treatment, plasmids used for HDTVI, mice used in study and time-points analyzed. **(B)** Representative gross images from AKT-NICD control show multiple and large tumors at 5 weeks while only few or no tumor seen in 5-Azacytidine-treated or *DN-TEAD* injected livers at the same time-point. Whereas widespread of *Akt-NICD*-driven cystic ICC nodules were observed in *Dnmt1* re-expressed livers in the setting of *TEAD* repression. **(C)** LW/BW ratio depicts comparable low tumor burden in 5-Azacytidine-treated, *DN-TEAD* or *DN-TEAD-Dnmt1* co-injected animals at 2 weeks. **(D)** Representative tiled image of HA-tag (AKT) staining showing near-complete abrogation of tumor by *TEAD* repression whereas widespread of cystic ICC nodules were detected in *Dnmt1* re-expressed livers. **(E)** IHC staining of squared region (D panel) showing HA^+^;V5 tag (*Dnmt1*)^+^;endogenous DNMT1^+^ ICC nodules in *Dnmt1* re-expressed liver (bottom). Likewise, few tumor observed in AKT-NICD *DN-TEAD* was V5^−^ but positive for HA and endogenous DNMT1 (top). **(F)** Representative gross images from *Akt-Dnmt1*-injected liver shows tumor burden within entire liver lobes while no gross tumor nodule was observed in *Dnmt1-Empty*-injected livers. **(G)** LW/BW ratio also depicts tumor development in *Akt-Dnmt1* livers with significantly higher LW/BW compared to *DNMT1-Empty* livers. **(I)** Representative IHC images of 5-week AKT-DNMT1 show HCC to be HNF4α^+^;panCK^−^ and few ICC components to be positive panCK but HNF4α^−^. Entire tumor burdens were V5 and DNMT1 positive regardless of tumor types. **(H)** While in DNMT1-Empty liver, no tumor was seen and all of V5^+^ cells were retained HC morphology without clonal expansion. Scale bars: 100μm; Error bar: standard error of the mean; *p<0.05; **p<0.01.

### *Dnmt1* re-expression restores abolished *Akt-NICD-driven-ICC* induced by TEAD inhibition

Next, to conclusively determine DNMT1 as a functional Notch-YAP/TEAD downstream effector in HC-driven ICC formation, we delivered plasmid expressing fulllength *Dnmt1* together with *Akt* and *NICD* along with DN-*TEAD* expression plasmid to rescue defective HC-to-ICC transformation in the setting of *TEAD* inhibition. As a control, *pT3-Ef1 α-Empty* was injected instead of *pT3-Ef1α-Dnmt1* plasmid. Remarkably, reexpression of *Dnmt1* restored *Akt-NICD-ICC* eliminated by *TEAD* inhibition. Gross and histologic observation clearly verified widespread HA^+^ ICC nodules by *Dnmt1* re-expression, whereas no or few nodules were observed in *DN-TEAD* livers (Fig.7B and 7D). LW/BW of *DN-TEAD-Dnmt1* liver was still comparable to *DN-TEAD* livers at this stage (Fig.7C) suggesting that DNMT1 partially rescues *Akt-NICD-ICC* formation blocked by TEAD repression. Microscopically, widespread ICC nodules in *DN-TEAD-Dnmt1-livers* were HA^+^;V5^+^;DNMT1 ^+^ whereas the majority of *Akt*-transduced cells (HA^*+*^) in *DN-TEAD*-livers were V5^−^;DNMT1^−^ with HC morphology indicating impaired HC-to-ICC conversion (Fig.7E). Interestingly, few ICC nodules in DN-*TEAD*-livers were V5^−^ but still strongly DNMT1^+^ addressing the indispensable role of DNMT1 in HC-to-ICC transformation (Fig.7E). Altogether, these data clearly demonstrate NICD-YAP/TEAD-DNMT1 axis essential for HC-driven cholangiocarcinogenesis.

### Co-expression of *Dnmt1* and *Akt* in HC provokes mixed HCC/ICC

Given the critical roles for DNMT1 in NICD-YAP-dependent HC-driven ICC development, we next tested if co-expression of *Dnmt1*, instead of *NICD* or *Yap*, along with *Akt* in HC is sufficient to induce tumor development. As controls, we injected *Dnmt1* or *pT3-Ef1a-Empty* plasmid. Surprisingly, co-expression of *Akt-Dnmt1* in HC induced liver cancer with significantly higher LW/BW at 5 weeks post HDTVI, while *Dnmt1* singular injection displayed normal LW/BW without any gross/microscopic tumor burden (Fig.7F and G). Microscopically, majority of *Dnmt1*-transduced cells (V5^+^;DNMT1^+^) were panCK^−^;HNF4α^+^ HCC-like tumors while a small number of V5^+^;DNMT1^+^;panCK^+^;HNF4α^−^ ICC nodules were also observed in the *Akt-Dnmt1* livers (Fig.7I). Altogether, co-expression of *Akt* and *Dnmt1* in HC was sufficient to provoke HCC-dominant mixed HCC/ICC in mouse.

## Discussion

ICC is among the most devastating tumors with less than a year survival from the time of diagnosis. Like other malignancies, varied response to drug candidates based on diverse molecular features, indicates that biomarker-based precise targeted therapy may be an ideal approach to treat this tumor. Along these lines, several novel compounds have been recently approved for use in ICC with clinically valid biomarkers, but so far, the overall impact on patient survival even on the stratified patients, has been limited (4–6). Molecular heterogeneity along with an acquired resistance to the selective chemotherapy, have been the major causes of tumor relapse and failure of the personalized approach. Thus, generation of clinically validated and biomarker-driven ICC models may be useful for delineating underlying cellular and molecular mechanisms and will be key to developing next-generation therapies against this deadly cancer.

Active cell fate transition between HC and BEC as well as activation of intermediate cell population (LPCs and ductular reaction) in diseased liver supports the concept that a subset of ICCs may be derived from HCs or LPCs. The plasticity of hepatic epithelial cells to reprogram into each other during hepatobiliary injury is now well recognized (20). While this is an important tier of repair, such reprogramming entails remodeling of chromatin structure and the epigenome, increasing temporal susceptibility to a stochastic event which may cumulatively lead to a higher risk of malignant transformation of a cell (38,39). Given the frequent observation of HC-to-BEC trans-differentiation in liver diseases that are known as major risk factors for ICC, identifying drivers of reprogramming is thus of high relevance.

We identified the presence of p-AKT as well as NICD targets YAP and SOX9 in two well known risk factors of ICC in patients. A subgroup of PSC and NASH patients showed increased expression of these HC-to-BEC reprogramming mediators in a subset of hepatocytes. Both PSC and NASH are associated with risk of ICC development (40,41). In conditions such as PSC where BECs are the afflicted cell type, upregulation of biliary markers in HCs may be a reparative response, whereas the upregulation of biliary markers in NASH may be an adaptive or dedifferentiation response to lipid accumulation in the hepatocyte and requires further mechanistic elucidation. Aberrant expression of these factors in subsets of hepatocytes may facilitate their transformation by altering their differentiation state and thus may be the first molecular hit needed for transformation, while additional mutagenic signals may be emanating from the adverse microenvironment of ongoing inflammation and ROS due to the disease process. While most CCA showed YAP and SOX9 expression since they are markers of BECs, the enrichment of p-AKT in ICC and mixed ICC-HCC suggests a unique role of AKT activation in cooperating with Notch, *Yap* and *Sox9* signaling in disease initiation and progression. Further studies are needed to address the role of AKT activation in this context. Suffice to say, that given the significant enrichment for p-AKT, SOX9, YAP and DNMT1 in HC from PSC liver, further studies are required to elucidate the clinical relevance of these as potential biomarkers for ICC development.

Although the cystic histology of ICC in the AN model is not commonly observed in human ICC, we believe this to be due to the technique to SB-HDTVI which delivers DNA into a subset of HCs located around zone-3 of a lobule. It is likely that these cystic patterns emerge due to spatial constraints faced by these ‘transfected’ cells as they proliferate under the “inside-out” oncogenic signals. In fact, molecular analysis of AN tumors showed significant genetic similarity to a subset of Notch-driven, proliferative subtype, and stemcell-like ICCs seen in patients. Transcriptomic analysis yielded DEGs which upon pathway analysis revealed several overlapping signaling pathways to human ICCs. Top DEGs were also applied to three independent human studies - GSE33327, GSE26566 and GSE76297 to generate unique signature which was then used for NTP analysis. NTP analysis provides class prediction with confidence, computed in each single patient’s gene-expression data using only a list of signature genes from the test dataset (42). We show that AKT-NICD model signature predicted ICC in 13% of cases in GSE33327, 12% in GSE26566 and 30% in GSE76297. Furthermore, most of these cases belonged to the “proliferation class” of ICC in all datasets as defined in an earlier study (21). In addition, in all three datasets, the group of human patients resembling the AKT-NICD murine model were enriched in NOTCH signaling (p<0.05) (25) and in 1/3 studies in the stem cell-like signature of human ICC (26).

We demonstrate YAP and SOX9 to be critical in most AKT-NICD-driven ICCs, although each exhibited overlapping and unique functions in tumorigenesis. We also observed high frequency of nuclear SOX9 and YAP in human CCAs, but interestingly also found a small subset of tumors exhibiting nuclear SOX9 or YAP singly. Since simultaneous ablation of *Yap* and *Sox9* prevented HC-derived ICC in the AKT-NICD model, these single-positive tumors may have distinct characteristics thus implying molecular mechanism for adaptive resistance against the targeting respective gene in ICC. In *Akt-NICD*-ICC formation, YAP, but not SOX9, appears to be essential for completion of HC-to-BEC reprogramming since its deletion resulted in incomplete repression of HNF4α. Indeed, YAP has been shown to be critical for biliary lineage commitment (15,43) as well as malignant HC-to-BEC trans-differentiation through the regulation of Notch-SOX9 cascade in the absence of Notch activity (27). However, in *Akt-NICD*-driven ICC development, we revealed a novel molecular mechanism: YAP regulates DNMT1 provoking HC-to-BEC conversion when Notch signaling is active, indicating the context-dependent diverse molecular mechanisms and crosstalk of these well studied biliary drivers in committing biliary fate in diseased liver.

There have been several studies demonstrating the critical roles for DNMT in liver cancer development and maintenance (31,44–46); most of studies showed that pharmacologic DNMT inhibition leads to hypomethylation of the promoters of various tumor suppressors such as *p16INK4a, FoxM1, TP53, PTEN*, and others (47), which typically induces death of malignant cells. At the same time, like other epigenetic regulators (20,48,49), DNMT1 plays critical roles in various cell fate switches including HC-to-BEC conversion (29–32,50). Likewise, we also found that DNMT1 inhibition impairs AN-mediated HC-to-BEC conversion, thereby completely abrogating HC-driven murine ICC development. Although two studies demonstrated that *FOXA2* and/or *SOX17* are downstream effectors of DNMT in BEC injury and ICC development (30,31), our RNA-seq data showed no significant difference in *Foxa2* and *Sox17* expression between AN-ICC and healthy liver (data not shown), suggesting DNMT1 mediates distinct molecular mechanisms in HC-driven ICC. Thus, together with high prevalence of DNMT1 in human ICC, it will be important to identify epigenetic targets of DNMT1 in the context of HC-driven ICC development through further studies.

## Materials and Methods

### Mouse husbandry and breeding

All animal care and experiments were performed in accordance with the Institutional Animal Care and Use Committee (IACUC) at the University of Pittsburgh. *Sox9^(fl/fl)^* and *Yap^(fl/fl)^* mice were purchased from Jackson Laboratories for breeding. All transgenic and KO mouse lines were maintained on the immunocompetent C57BL/6 genetic background. All animals ranged from 8-12 weeks in age for analysis and were from either sex.

### Patient data

Study approval for all human tissue samples was obtained from the University of Pittsburgh (IRB# STUDY19070068). All samples were provided by the Pittsburgh Liver Research Center’s (PLRC’s) Clinical Biospecimen Repository and Processing Core (CBPRC), supported by P30DK120531. Tissue sections were obtained from archival formalin-fixed paraffin-embedded tissue blocks from 6 patients with healthy liver who underwent biopsy secondary to colon adenocarcinoma, 10 patients with PSC, and 10 patients with NASH (Supplementary Table 1). Sections were stained manually using antibody against p-AKT-S473 (Cell Signaling), SOX9 (EMD Millipore) and YAP (Cell Signaling) as described in IHC sections. The staining was scored as 0 (negative, 0-5 positives hepatocytes observed in a whole section); 1 (positive, <20% hepatocytes positive within a section); and 2 (positive, 20-50% of hepatocytes positive within a section. Scores of 0 are considered negative (NEG), and 1 and 2 are considered positive (POS).

TMAs were constructed from archival formalin-fixed paraffin-embedded tissue blocks from 108 cholangiocarcinoma patients seen at the University of Pittsburgh Medical Center and were also obtained from PLRC’s CBPRC supported by P30DK120531. All tumor hematoxylin and eosin (H&E) stained slides were reviewed, and representative areas were carefully selected for tissue microarray construction. Two, random 1.0 mm-sized cores were punched from each patient’s tumor and harvested into recipient blocks. The demographics and additional information of these cases are included in Supplementary Table 2. The TMA were stained manually using antibody against p-AKT-S473 (Cell Signaling), SOX9 (EMD Millipore), YAP (Cell Signaling) and DNMT1 (Abcam) as described in IHC sections. Whole slide image capture of the tissue microarray was acquired using the Aperio XT slide scanner (Aperio Technologies). The staining was evaluated and scored by anatomic pathologist (A.S.). Staining for SOX9 and YAP was scored either as 0 (negative), 1 + (mostly cytoplasmic staining or very weak staining in CC tumor cells), or 2+ (strong positive nuclear staining in CC tumor cells). Staining for pAKT was scored either as 0 (negative), 1+ (weak positive cytoplasmic staining), 2+ (moderately strong positive cytoplasmic staining in majority of CC tumor cells) or 3+ (strong positive cytoplasmic staining in majority of CC tumor cells). For all 4 markers, the scores for different tissue sections from each patient were averaged to get a single score per patient (Supplementary Table 2). Mean scores greater than or equal to 1.5 were considered “HIGH” and mean scores less than 1.5 were considered “LOW/NEGATIVE”.

Representative images are included in Supplementary Fig. S1D and all scores for each individual section are included in Supplementary Table 2.

**Please see Online Supplement for Additional Methods**

## Authors’ Contributions

<u>Conception and design:</u> SK, SPM

<u>Development of methodology:</u> SH, AB, SK, JT

<u>Acquisition of data (provided animals, acquired and managed patients, provided facilities, etc.):</u> MH, LM, SH, SS, MP, KNB, MO, RR, AS, SPM

<u>Analysis and interpretation of data (e.g., statistical analysis, biostatistics, computational analysis):</u> SK, SH, DS, LM, SL, MO, AS, SPM

<u>Writing, review, and/or revision of the manuscript:</u> SK, LM, SPM

<u>Administrative, technical, or material support (i.e., reporting or organizing data, constructing databases):</u> SH, SS, DS, MP, AB

<u>Study supervision:</u> SK, SPM

