## Supplementary material for "NOTCH-YAP1/TEAD-DNMT1 axis regulates hepatocyte reprogramming into intrahepatic cholangiocarcinoma": Online Supplement

1. **SUPPLEMENTARY METHODS (WITH REFERENCES)**
2. **SUPPLEMENTARY TABLE**
3. **SUPPLEMENTARY FIGURES AND LEGENDS**

### 1. SUPPLEMENTARY METHODS

**Animal Models of Intrahepatic Cholangiocarcinoma.** The constructs used for mouse SB-HDTVI, including *pT3-EF1α, pT3-EF1α- myrAkt-HA* (mouse), *pT3-EF1α-Myc-N1ICD* (mouse), *pT3-EF1α-YapS127A* (human), *pCMV-empty*, *pCMV-Cre, pT3-EF1α-Sox9* (mouse), *pT3-EF1α-HA-myrAkt-sh-Luciferase, pT3-EF1α-HA-myrAkt-Sh-Yap, pT3-EF1α-myrAkt-HA-Sh-Dnmt1 (mouse), pT3-EF1α-Dnmt1-V5* (mouse), *DN-Tead-V5, Fbxw7, Kras, sh-p53* and *pCMV-sleeping beauty transposase* (SB) were generated or have been described elsewhere (1-3). All the plasmids used for *in vivo* experiments were purified using the Endotoxin Free Maxi Prep kit (Sigma-Aldrich). Six-to-eight weeks old mice were randomized into groups and subjected to the sleeping beauty transposon-transposase and hydrodynamic tail vein (SB-HTVI) protocol as described previously (1). Briefly, 10 μg *pT3-EF1α-myrAkt-HA*, 20 μg of *pT3-EF1α-Myc-N1ICD* and μg mg of *pCMV-Cre (or pCMV-empty)*, or 10 μg *pT3-EF1α-myrAkt-HA* and 20 μg of *pT3-EF1α-Sox9*, or μg mg of *pT3-EF1α-myrAkt-HA-sh-Luciferase*, 20 μg of *pT3-EF1α-Myc-N1ICD, or* 10 μg of *pT3-EF1α-myrAkt-HA-Sh-Yap*, 20 μg of *pT3-EF1α-Myc-N1ICD*, or 10 μg of *pT3-EF1α-myrAkt-HA-Sh-Dnmt1*, 20 μg of *pT3-EF1α-Myc-N1ICD, or* 10 μg of *pT3-EF1α-myrAkt-HA*, 20 μg of *pT3-EF1α-Myc-N1ICD,* 60 μg *of pT3-EF5α-Dn-TEAD, or* 20 μg of *pT3-EF1α-Myc-N1ICD, 60* μg *of pT3-EF5α-Dn-TEAD, or* 10 μg of *pT3-EF1α-myrAkt-HA-sh-Yap*, 20 μg of *pT3-EF1α-Myc-N1ICD*, 10 μg and 40 μg of *pT3-EF1α-Dnmt1-V5, or* 10 μg of *pT3-EF1α-myrAkt-HA*, 20 μg of *pT3-EF1α-Myc-N1ICD, 60* μg *of pT3-EF5α-Dn-TEAD, 40* μg *of pT3-EF1α-Dnmt-V5,* or 10 μg of *pT3-EF1α-myrAkt-HA-sh-Luciferase*, 20 μg of *pT3-EF1α-Myc-N1ICD* and 40 μg of *pCMV-empty*, or 10 μg of *pT3-EF1α-myrAkt-HA-sh-Yap*, 20 μg of *pT3-EF1α-Myc-N1ICD* and 40 μg of *pCMV-Cre*, or 10 μg of *pT3-EF1α-*

*myrAkt-HA*, 20 μg of *Fbxw7ΔF*, or 10 μg of *Kras*, 40 μg of *sh-p53* along with the transposase in a ratio of 25:1 were diluted in 2 ml of normal saline (0.9% NaCl), filtered through 0.22μm filter (Millipore), and hydrodynamically injected into the lateral tail vein of mice. All animals were sacrificed between 2-5 weeks of plasmids injections unless otherwise indicated.

**5-Azacytidine Treatment.** 5-Azacytidine (MCE, HY-10586) was dissolved in DMSO as a 100 mg/ml stock and stored at -80°C in aliquots. Mice were i.p. injected with either 1 mg/kg 5-Azacytidine dissolved in 0.9% saline and 0.9% saline alone as a control.

**Immunohistochemistry (IHC).** Mouse liver tissues were fixed for 48 h in 10% neutralized formalin (Fisher Chemicals), transferred into 70% ethanol and then dehydrated and embedded in paraffin. For IHC, formalin-fixed sections were deparaffinized in graded xylene and ethanol and rinsed in PBS. For antigen retrieval, samples were microwaved for 12 min in pH 6.0 sodium citrate buffer (HA-tag, Myc-tag, panCK, SOX9) or pH 9.0 Tris-EDTA buffer (p-AKT) or DAKO (V5-tag), or were pressure cooked for 20 min in pH 9.0 Tris-EDTA buffer (YAP and HNF4α), or were autoclaved for 1 h in pH 9.0 Tris-EDTA buffer (DNMT1). After cooling, samples were placed in 3% $H_2O_2$ (Fisher Chemicals) for 10 min to quench endogenous peroxide activity. After washing with PBS, slides were blocked with Super Block (ScyTek Laboratories) for 10 min. Sections were incubated for overnight at 4C with the primary antibodies (Supplementary Table 3). Sections were then incubated with species-specific biotinylated secondary antibodies (EMD Millipore, Supplementary Table 3) for 1 h, at room temperature. Sections were incubated with

Vectastain ABC Elite kit (Vector Laboratories) and signal was detected with DAB Peroxidase Substrate Kit (Vector Laboratories) followed by quenching in distilled water for 5 min. Slides were counterstained with hematoxylin (ThermoFisher Scientific), dehydrated to xylene (Fisher Chemicals) and coverslips applied with Cytoseal XYL (ThermoFisher Scientific).

**Immunofluorescence.** Paraffin embedded liver sections (5 μm thick) were deparaffinized using xylene (Fisher Chemicals) and rehydrated by incubating the slices in ethanol (100% and 95% v/v, each 3x5 min) and washed in PBS. Heat-induced epitope retrieval was performed for 20 min using a pressure cooker with pH 6.0 sodium citrate buffer. Sections were washed in PBS, permeabilized for 5 minutes with PBS/0.3% Triton X and blocked with PBS/0.3% Triton X/10% bovine serum albumin (BSA) for 45 minutes at room temperature. Sections were incubated with primary antibodies in PBS/0.3% Triton X/10% BSA overnight at 4C. At the end of the incubation, sections were washed thrice and incubated with fluorochrome-conjugated secondary antibodies in PBS/0.3% Triton X/10% BSA for 1h at room temperature, then washed in PBS/0.1% Triton X 3 times. Liver sections were mounted using Prolong Gold Antifade w/DAPI (Invitrogen) and pictures were acquired using LSM700 confocal microscope the and Zen Software (Zeiss).

**RNA-Seq Analysis.** For each group (wild type or WT, AKT-NICD), three mouse liver samples were processed for RNA-seq analysis. RNA samples from livers were used to generate library using TruSeq kit from Illumina (4,5). We used in-house HiSeq2500 platform and sequenced 200 million reads to accurately quantify genes & transcripts (6).

Raw sequencing data was analyzed by FastQC for quality control (7). Low quality reads or adapter sequences were trimmed out by Trimmomatic (8). After pre-processing, sequenced reads were aligned to mouse reference genome mm10 by HISAT2 aligner (9). Read counts for each gene were then quantified by HTSeq (10). All the pipelines were run by default parameter settings.

Differential expression (DE) analysis was performed to compare WT versus AKT-NICD. Based on the read counts, DE tests were performed by R package ‘DEseq2’ (11) and top DE genes were selected by absolute fold-change greater than 1 and FDR=0.05. These DE genes were then used as input for Ingenuity Pathway Analysis (IPA)® to call pathways that were significantly enriched with FDR=0.1.

**Bioinformatic comparison between mouse and human models.** To further investigate how the mouse models may mimic a subset of human ICC or human HCC, public human transcript data were collected to compare with the gene signature obtained from the mouse models. Three human datasets were downloaded from NCBI Gene Expression Omnibus (GEO) database: GSE33327 (12), GSE35306 (13) and GSE76297 (14). For HCC, LIHC TCGA database was assessed similarly. For the microarray data, samples were first pre-processed by quantile normalization to scale the expression in the same level (15). Probe-based intensities were then mapped to gene-based expression. If multiple probes were annotated with the same gene, only the probe with the largest interquartile range will be utilized as representative of that gene. After pre-processing, gene expression data were analyzed by R package ‘limma’ test (16) and top differentially expressed genes were selected by absolute fold-change greater than 1 and FDR=0.05 (same criteria as the mouse model). Top DE genes from human study were further used to detect significantly enriched pathways by IPA® software with FDR=0.1.

To test the molecular similarity between our mouse AKT-NICD ICC model and the human studies, three publicly available CCA human datasets (GSE33327, GSE26566 and GSE76297) were analyzed by three comparisons. (1) Pathway enrichment analysis. Top enriched pathways detected by mouse model and human studies were detected independently and compared. (2) Gene signature analysis. Top DE genes from mouse models were converted to human homologous genes by Mouse Genome Database (MGD) (17). These genes were then applied into the human studies to check their expression signatures. (3) Signature prediction. Positive prediction of gene signatures tested in this study has been calculated by Nearest Template Prediction (Gene Pattern module), (18) as previously described (19). For ICC studies, each patient was also assigned to either the Proliferation or Inflammation class based on the results of unsupervised clustering as previously described (12). Additional signatures assessed were for Notch activation (20) and hepatic stem-cell like group of ICC patients (21). Any significant correlation to a subclass was noted by p value ($p<0.05$).

**ChIP-Seq Data Analysis.** Public ChIP-seq data were obtained from Gene expression omnibus (GEO) database with accession ID GSE107860 (22). Library for TEAD4 binding to TetO-YAP mice were specifically analyzed. Quality control was first performed on the raw sequencing data using FastQC software and low-quality reads and adapter sequences were removed by tool Trimmomatic (8). Then the surviving reads were aligned to mouse reference genome mm10 by Burrows-Wheeler Aligner (BWA) software (23) and duplicate reads were marked by Picard online tool. Eventually, peaks were called by Model-based analysis of ChIP-Seq (MACS) tool (24) to detect local enriched TEAD4 binding sites. Selected binding regions were annotated to mouse genes for downstream analysis.

**Searching for Transcription Factor Binding Site.** Validated binding motives for transcription factor TEAD2, TEAD3 and TEAD4 were initially obtained from *TRANSFAC* database. Then tool FIMO [FIMO] were applied to scan the 5,000 bp upstream sequence of *Dnmt1* promoter region for respective motives. Significant matching sequences ($p \leq 0.0001$) and their corresponding motives were displayed.

**Statistical Analysis.** For all mouse experiments, sample size was pre-determined based on previous literature describing SB-HDTVI-mediated liver carcinogenesis (1). Accordingly, littermates were randomized into groups for HDTVI and managed throughout the course of treatment in a non-blinded manner. All subsequent molecular, immunohistochemical, and immunofluorescence analysis was performed in a blinded manner. All confidence intervals shown on the bar plots are presented as mean ± standard error of mean (SEM). Differences in mean values of liver volume and LW/BW ratio were analyzed by one-way ANOVA assuming normal Gaussian distribution with Geisser Greenhouse posttest correction. For patient data, Fisher’s exact test (two-sided) with post hoc pairwise comparison was utilized to assess statistical significance. p<0.05

was considered significant (*), $p<0.01$ was considered highly significant (**), $p<0.005$ was considered extremely significant (***), and so on. All statistical analysis on patient samples has been included in the results section and respective p-values were included in the pertinent text and figure legends. All statistics were performed using GraphPad Prism 8.0 (GraphPad Software) or R software.

### 2. Supplementary Table

#### Supplementary Table 1: Summary of Patients with PSC and NASH used in the study.

| Patient Record Number | Age | Sex | SOX9 | pAKT S473 | YAP | Hepatocyte DNMT1 | | Disease |
|---|---|---|---|---|---|---|---|---|
| | | | | | | C | N | |
| PHR17-187 | 44 | F | <20% | NEG | 20-50% | 100% | NEG | PSC |
| PHR17-315 | 39 | M | NEG | NEG | <20% | 100% | NEG | PSC |
| PHR17-318 | 32 | M | 20-50% | NEG | <20% | 50% | 20-50% | PSC |
| PHR17-340 | 47 | F | 20-50% | <20% | 20-50% | 100% | 20-50% | PSC |
| PHR17-471 | 44 | F | <20% | <20% | NEG | 100% | <20% | PSC |
| PHS18-35 | 35 | M | <20% | <20% | NEG | 0% | NEG | PSC |
| PHR19-64 | 24 | M | <20% | 20-50% | 20-50% | 100% | <20% | PSC |
| PHR19-67 | 32 | F | NEG | NEG | <20% | | | PSC |
| PHR19-74 | 50 | M | <20% | 20-50% | <20% | 100% | <20% | PSC |
| PHR19-80 | 61 | F | NEG | 20-50% | <20% | 100% | NEG | PSC |
| PHR16-107 | 67 | M | <20% | <20% | <20% | 0% | 0 | NASH |
| PHR17-50 | 73 | F | <20% | NEG | <20% | 0% | NEG | NASH |
| PHR17-114 | 53 | F | 20-50% | 20-50% | NEG | | | NASH |
| PHR17-235 | 63 | M | 20-50% | 20-50% | NEG | 0% | NEG | NASH |
| PHR17-238 | 65 | M | <20% | 20-50% | NEG | 50% | NEG | NASH |
| PHR17-247 | 61 | M | <20% | NEG | <20% | | | NASH |
| PHR17-299 | 56 | F | <20% | <20% | <20% | <20% | 0 | NASH |
| PHR18-25 | 68 | F | 20-50% | 20-50% | <20% | 50% | <20% | NASH |
| PHR18-219 | 62 | F | <20% | NEG | <20% | <20% | <20% | NASH |
| PHR19-107 | 78 | F | <20% | <20% | <20% | 100% | NEG | NASH |
| NHL1029 | | | NEG | NA | NEG | | | Healthy control liver |
| 16-5566 | 54 | F | NEG | NEG | NEG | | | Healthy control liver (underlying colon adenocarcinoma) |

| | | | | | | | | |
|---|---|---|---|---|---|---|---|---|
| 12-9718 | 60 | M | NEG | NEG | NEG | | | Healthy control liver (underlying colon adenocarcinoma) |
| 17-9693 | 49 | F | NEG | NEG | NEG | | | Healthy control liver (underlying colon adenocarcinoma) |
| 13-15397 | 54 | F | NEG | NEG | 20-50% | | | Healthy control liver (underlying colon adenocarcinoma) |
| 14-2191 | 36 | M | NEG | NEG | NEG | | | Healthy control liver (underlying colon adenocarcinoma) |
| PHS18-34295 | | | | | | 0 | NEG | Healthy control liver |
| PHS18-11592 | | | | | | 0 | NEG | Healthy control liver |
| PHS18-5904 | | | | | | 0 | NEG | Healthy control liver |
| PHS16-40535 | | | | | | 0 | NEG | Healthy control liver |
| PHS16-32066 | | | | | | 0 | NEG | Healthy control liver |

**Abbreviations: C-**cytoplasmic; **N-** nuclear; **PSC-** Primary sclerosing cholangitis; **NASH-** Non alcoholic steatohepatitis

**Supplementary Table 2: Attached independently as an excel spreadsheet**

**Supplementary Table 3: Antibody List used in the study**

**Primary Antibodies:**

| **Target** | **Species** | **Dilution** | **Source** | **Catalog Number** |
|---|---|---|---|---|
| HA-tag | Mouse | 1:50 (IHC) | Cell Signaling | CS2367 |
| Myc-tag | Rabbit | 1:100 (IHC) | Maine Medical Center Research Institute (mmcri) | Vli01 |
| Pan-CYTOKERATIN (panCK) | Rabbit | 1:200 (IHC) | Dako | Z0622 |
| p-AKT | Rabbit | 1:100 (IHC) | Cell Signaling | CS4060 |
| SOX9 | Rabbit | 1:200 (IF)<br>1:2000 (IHC) | EMD Millipore | Ab5535 |
| YAP | Rabbit | 1:100 (IF and IHC) | Cell Signaling | CS14074 |
| HNF4$\alpha$ | Rabbit | 1:100 (IHC) | Cell Signaling | CS3113 |
| PCNA | Mouse | 1:200 (IF) | Santa Cruz | sc-56 |
| CK19 | Rat | 1:10 (IF) | DSHB | TROMA III |
| DNMT1 | Rabbit | 1:100 (IHC) | Abcam | ab188453 |

**Secondary Antibodies:**

| **Secondary Antibody** | **Species** | **Source** | **Catalog Number** |
|---|---|---|---|
| Donkey anti-Rabbit IgG Biotin | Donkey | EMD Millipore | AP182B |
| Goat anti-Mouse IgG (H+L) Biotin | Goat | EMD Millipore | AP181B |
| Alexa-Fluor 555 Donkey anti-Rabbit IgG (H+L) | Donkey | Invitrogen | A31572 |
| Alexa-Fluor 488 Donkey anti-Rat IgG (H+L) | Donkey | Invitrogen | A21208 |
| Alexa-Fluor 647 Goat anti-Mouse IgG (H+L) | Goat | Invitrogen | A21236 |

### 3. Supplementary Figures and Legends

#### Figure S1

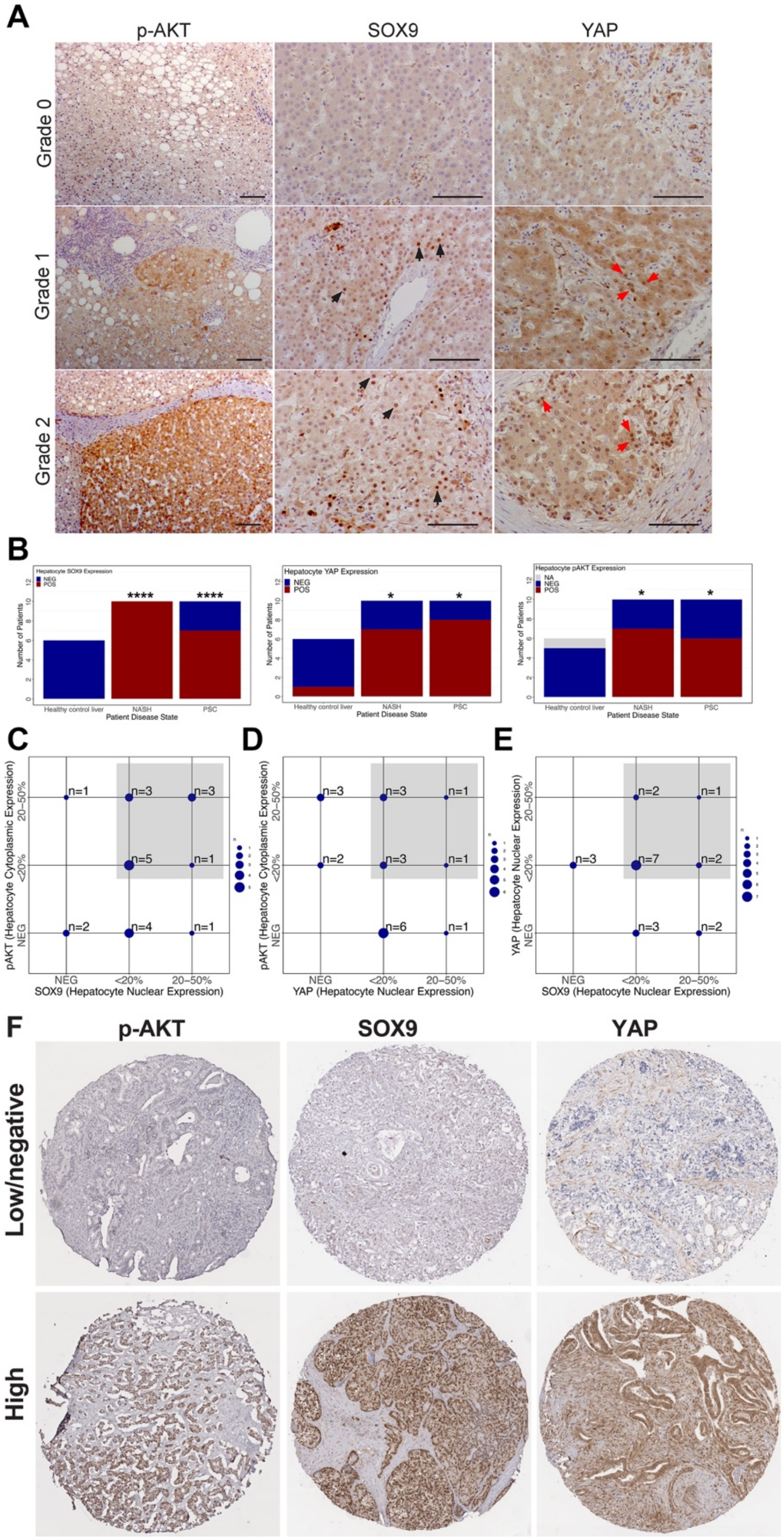

**Figure S1. Activation of p-AKT, YAP and SOX9 in human liver diseases with higher risk for the development of ICC**

(**A**) Representative human liver section images used for arbitrary scoring for p-AKT, SOX9 and YAP expression by IHC. Grade 0, no expression; grade 1< 20%; grade 2<50% (Grade 1 and 2 are considered positive). (**B)** Comparison of the number of patients with positive expression of SOX9, YAP, or p-AKT in HCs shows an enrichment in positive HCs among PSC and NASH patients vs healthy controls. **(C-E)** Overlap in the expression of YAP, SOX9, and p-AKT among patients with PSC and NASH shows most patients have at least 2 of 3 markers upregulated. (**F**) Representative human CCA TMA images used for arbitrary scoring for p-AKT, SOX9 and YAP expression by IHC (High vs Low/negative). Scale bars:100 μm. *p<0.05; ****p< 0.0001.

**Figure S2**

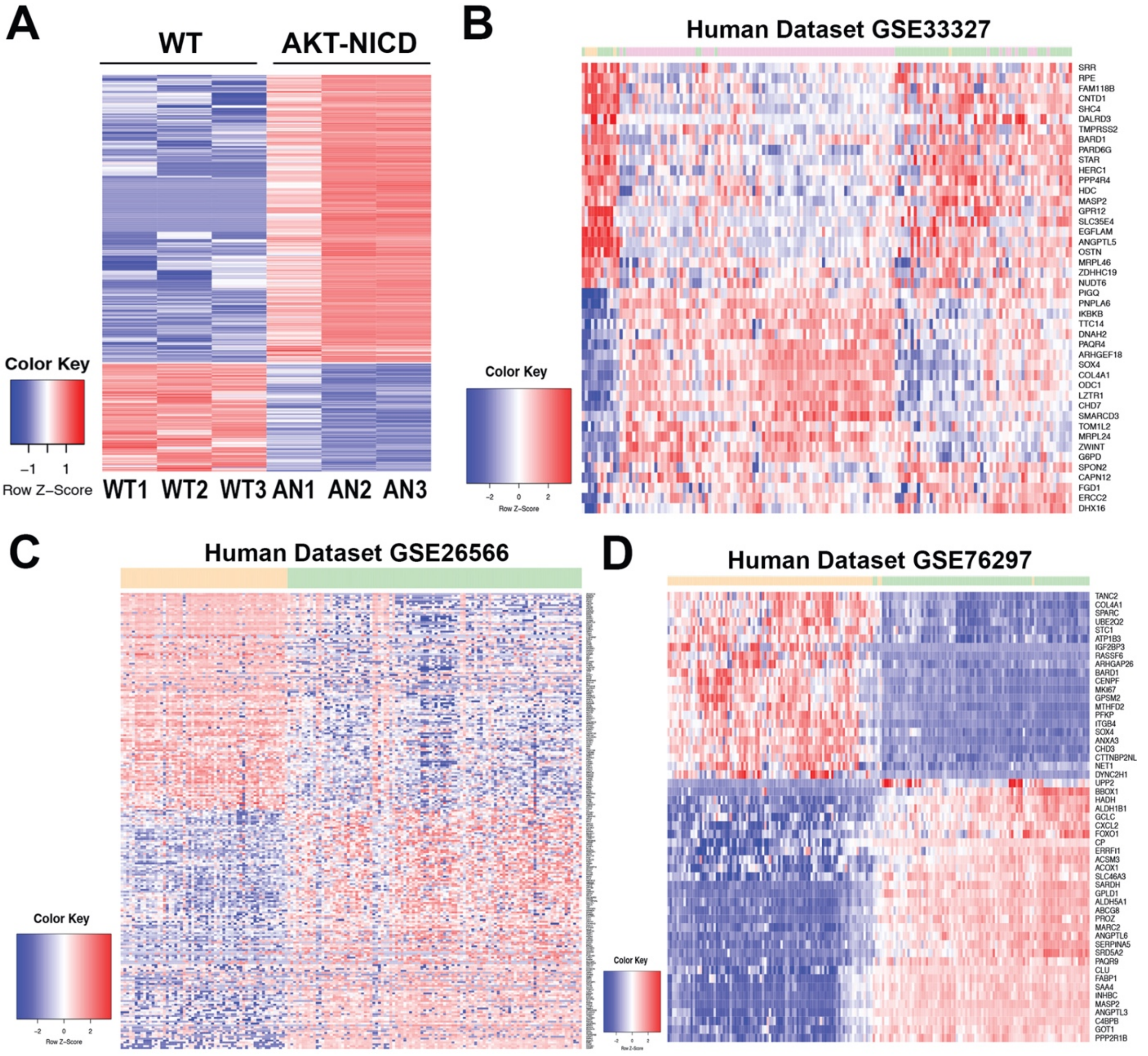
A
WT
AKT-NICD
Color Key
−1 1
Row Z−Score
WT1 WT2 WT3 AN1 AN2 AN3
B
Human Dataset GSE33327
SRR
RPE
FAM118B
CNTD1
SHC4
DALRD3
TMPRSS2
BARD1
PARD6G
STAR
HERC1
PPP4R4
HDC
MASP2
GPR12
SLC35E4
EGFLAM
ANGPTL5
OSTN
MRPL46
ZDHHC19
NUDT6
PIGQ
PNPLA6
IKBKB
TTC14
DNAH2
PAQR4
ARHGEF18
SOX4
COL4A1
ODC1
LZTR1
CHD7
SMARCD3
TOM1L2
MRPL24
ZWINT
G6PD
SPON2
CAPN12
FGD1
ERCC2
DHX16
Color Key
-2 0 2
Row Z-Score
C
Human Dataset GSE26566
Color Key
-2 0 2
Row Z-Score
D
Human Dataset GSE76297
TANC2
COL4A1
SPARC
UBE2Q2
STC1
ATP1B3
IGF2BP3
RASSF6
ARHGAP26
BARD1
CENPF
MKI67
GPSM2
MTHFD2
PFKP
ITGB4
SOX4
ANXA3
CHD3
CTTNBP2NL
NET1
DYNC2H1
UPP2
BBOX1
HADH
ALDH1B1
GCLC
CXCL2
FOXO1
CP
ERRFI1
ACSM3
ACOX1
SLC46A3
SARDH
GPLD1
ALDH5A1
ABCG8
PROZ
MARC2
ANGPTL6
SERPINA5
SRD5A2
PAQR9
CLU
FABP1
SAA4
INHBC
MASP2
ANGPTL3
C4BPB
GOT1
PPP2R1B
Color Key
-2 0 2
Row Z-Score

**Figure S2. RNA-sequencing analysis of AKT-NICD-driven murine ICC and comparison with human ICC studies.**

**(A)** Heatmap for the differentially expressed genes comparing wild-type mouse and AKT+NICD mouse. **(B)** Heatmap of gene signatures in human GSE33327 study that are selected by mouse model (WT VS AKT+NICD). **(C)** Heatmap of gene signatures in human GSE26566 study that are selected by mouse model (WT VS AKT+NICD). **(D)** Heatmap of gene signatures in human GSE76297 study that are selected by mouse model (WT VS AKT+NICD).

### Figure S3

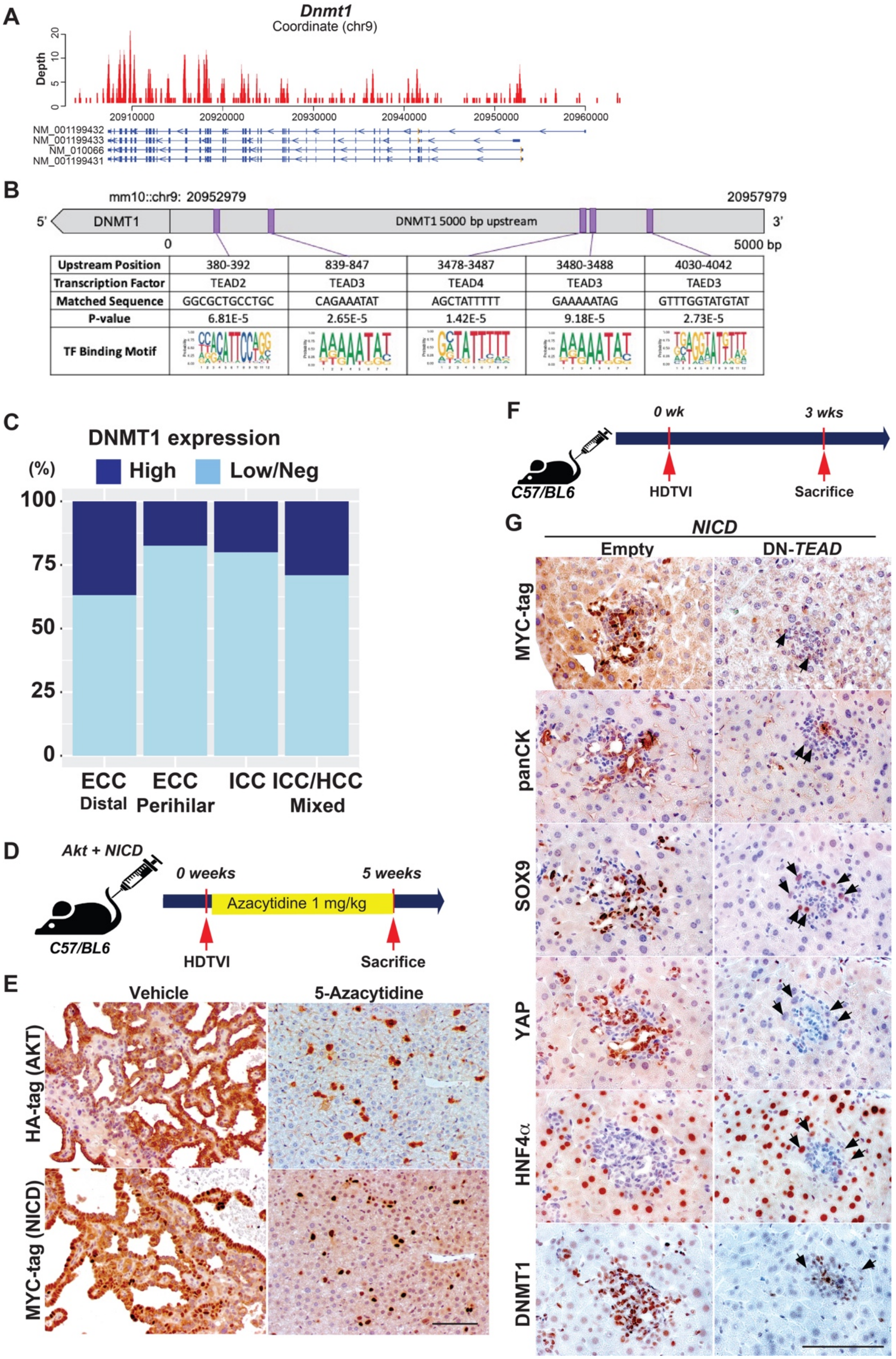
A
Dnmt1
Coordinate (chr9)
Depth
NM_001199432
NM_001199433
NM_010066
NM_001199431
B
mm10::chr9: 20952979
20957979
5'
DNMT1
DNMT1 5000 bp upstream
3'
0
5000 bp
Upstream Position | 380-392 | 839-847 | 3478-3487 | 3480-3488 | 4030-4042
Transcription Factor | TEAD2 | TEAD3 | TEAD4 | TEAD3 | TAED3
Matched Sequence | GGCGCTGCCTGC | CAGAAATAT | AGCTATTTTT | GAAAAATAG | GTTTGGTATGTAT
P-value | 6.81E-5 | 2.65E-5 | 1.42E-5 | 9.18E-5 | 2.73E-5
TF Binding Motif
C
DNMT1 expression
(%)
High
Low/Neg
ECC Distal
ECC Perihilar
ICC
ICC/HCC Mixed
D
Akt + NICD
0 weeks
5 weeks
Azacytidine 1 mg/kg
C57/BL6
HDTVI
Sacrifice
E
Vehicle
5-Azacytidine
HA-tag (AKT)
MYC-tag (NICD)
F
0 wk
3 wks
C57/BL6
HDTVI
Sacrifice
G
NICD
Empty
DN-TEAD
MYC-tag
panCK
SOX9
YAP
HNF4α
DNMT1

**Figure S3. Supporting data for Figure 5 and 6. (A)** ChIP-seq data for TEAD4 binding sites on *Dnmt1* genomic regions are collected form GEO GSM2882182. **(B)** Predicted TEAD2/TEAD3/TEAD4 binding sites located at 5,000 bp upstream from *Dnmt1* transcription start site. **(C)** Stratifying DNMT1 staining by CCA subtypes shows overall 27.8% prevalence without any specific enrichment among CCA subtypes. **(D and F)** Experimental design illustrating 5-Azacytidine treatment, plasmids used for HDTVI, mice used in study and time-points analyzed. **(E)** Representative IHC images for HA-tag (AKT), MYC-tag (NICD), and panCK showing dramatic tumor development in AKT-NICD WT livers and normal histology in 5-Azacytidine-treated livers at 5 weeks post HDTVI. **(G)** Representative IHC staining of *NICD-Empty* or *NICD*-DN-*TEAD* injected livers showing NICD-transfected cells (Myc tag$^+$) with intact HC-to-ICC reprogramming with loss of HNF4$\alpha$ and gain of YAP, panCK and DNMT1. DN-*TEAD* livers showing *NICD*-transfected cells with defective HC-to-BEC reprogramming with continued HNF4$\alpha$ staining and absent of DNMT1 in YAP$^-$ HCs (black arrows). Scale bars:100 μm.

### Figure S4

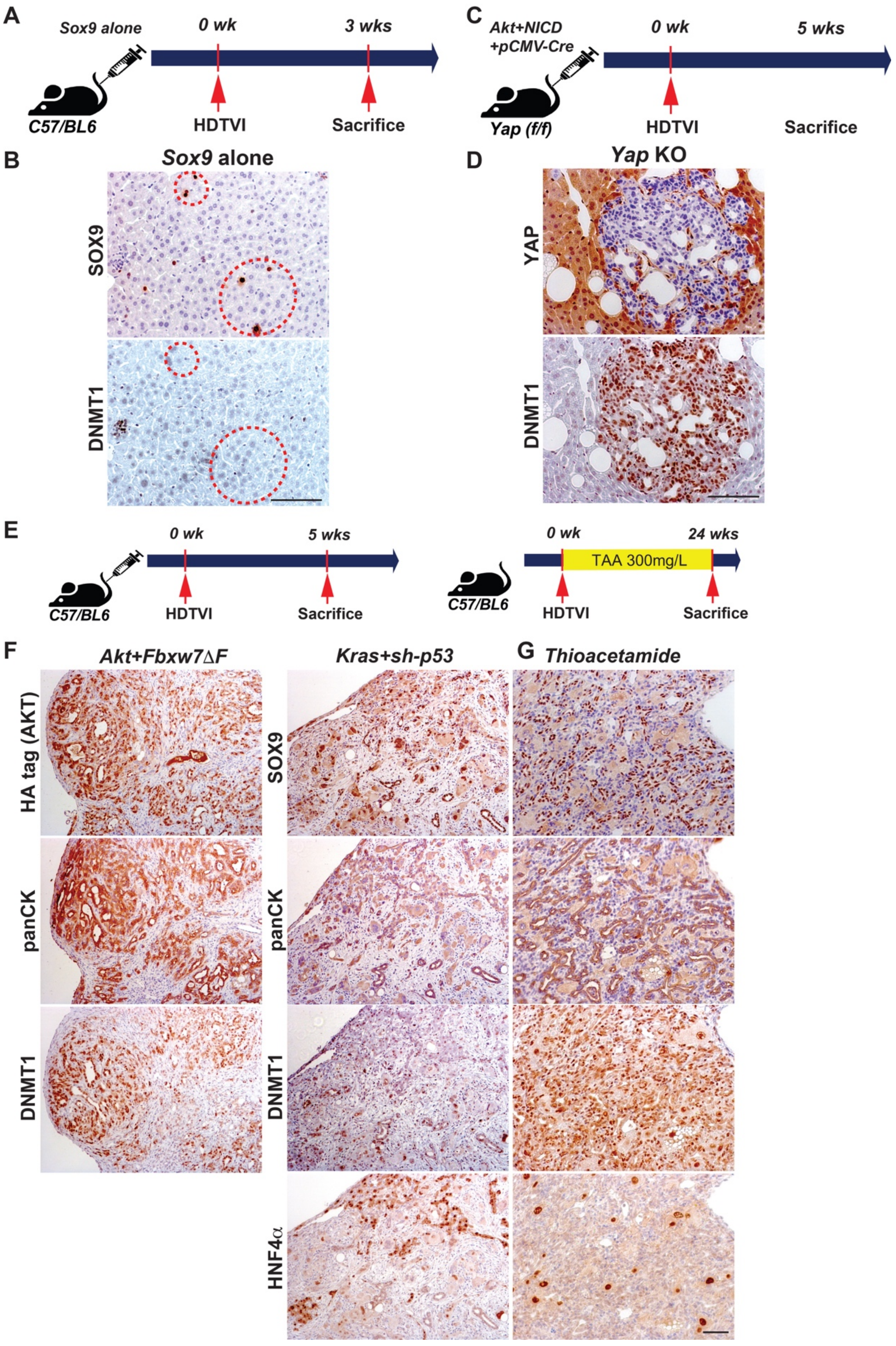
A
Sox9 alone
0 wk
3 wks
C57/BL6
HDTVI
Sacrifice
C
Akt+NICD
+pCMV-Cre
0 wk
5 wks
Yap (f/f)
HDTVI
Sacrifice
B
Sox9 alone
SOX9
DNMT1
D
Yap KO
YAP
DNMT1
E
0 wk
5 wks
C57/BL6
HDTVI
Sacrifice
0 wk
24 wks
TAA 300mg/L
C57/BL6
HDTVI
Sacrifice
F
Akt+Fbxw7ΔF
Kras+sh-p53
G
Thioacetamide
HA tag (AKT)
panCK
DNMT1
SOX9
panCK
DNMT1
HNF4α

**Figure S4. DNMT1 expression pattern in genetically modified HC-driven ICC models. (A, and C)** Experimental design illustrating plasmids used for HDTVI, mice used in study and time-points analyzed. **(B)** Representative IHC staining of *Sox9* expression plasmid-injected liver showing *Sox9* singular-transduced cells does not induce DNMT1 expression in HC. **(D)** IHC staining for *Akt-NICD-Yap* KO ICC at 5 weeks post HDTVI showing strong DNMT1 expression. **(E)** Experimental design illustrating thioacetamide treatment, HDTVI, mice used in study and time-points analyzed. **(F)** Representative IHC staining of liver from *Akt-Fbxw7ΔF* or *KRAS*-sh-*p53*-driven ICC shows strong DNMT1 expression. **(G)** Representative IHC staining of liver from TAA-administered mice at 24 weeks. IHC staining indicates panCK$^{+}$;SOX9$^{+}$;HNF4$\alpha^{-}$;DNMT1$^{+}$ ICC with biliary morphology.
